# Hominin-specific NOTCH2 paralogs expand human cortical neurogenesis through regulation of Delta/Notch interactions

**DOI:** 10.1101/221358

**Authors:** Ikuo K. Suzuki, David Gacquer, Roxane Van Heurck, Devesh Kumar, Marta Wojno, Angéline Bilheu, Adèle Herpoel, Julian Chéron, Franck Polleux, Vincent Detours, Pierre Vanderhaeghen

## Abstract

The human cerebral cortex has undergone rapid expansion and increased complexity during recent evolution. Hominid-specific gene duplications represent a major driving force of evolution, but their impact on human brain evolution remains unclear. Using tailored RNA sequencing (RNAseq), we profiled the spatial and temporal expression of Hominid-specific duplicated (HS) genes in the human fetal cortex, leading to the identification of a repertoire of 36 HS genes displaying robust and dynamic patterns during cortical neurogenesis. Among these we focused on NOTCH2NL, previously uncharacterized HS paralogs of NOTCH2. NOTCH2NL promote the clonal expansion of human cortical progenitors by increasing self-renewal, ultimately leading to higher neuronal output. NOTCH2NL function by activating the Notch pathway, through inhibition of Delta/Notch interactions. Our study uncovers a large repertoire of recently evolved genes linking genomic evolution to human brain development, and reveals how hominin-specific NOTCH paralogs may have contributed to the expansion of the human cortex.

## Introduction

The number and diversity of neurons is at the core of the unique computational capabilities of the human cerebral cortex. The neocortex in particular is arguably the most divergent brain structure among mammals. It has undergone a rapid and considerable increase in relative size and complexity over the last few millions of years of hominid evolution, with significant impact on the acquisition of higher functions in human species (Hill and Walsh, 2005; Lui et al., 2011; Rakic, 2009; Sousa et al., 2017). As the enlargement of the surface and thickness of the human cortex is largely due to increased number of cortical neurons (Amadio and Walsh, 2006; Rakic, 2009; Sousa et al., 2017), many of the species-specific features of the human cortex are likely linked to divergence in the mechanisms underlying the generation, specification, and differentiation of cortical neurons, a process referred to as cortical neurogenesis (Bystron et al., 2008; Fish et al., 2008; Geschwind and Rakic, 2013; Lui et al., 2011; Rakic, 2009).

While cortical neurogenesis appears to be well conserved among mammals, a number of divergent features have been identified, many of which are linked to larger expansion capacity and increased self-renewal potential of cortical progenitors, thereby allowing them to generate a higher number of neurons in the human brain (Fietz et al., 2010b; Fish et al., 2008; Hansen et al., 2010a; Kriegstein et al., 2006; Lui et al., 2011). Radial glial cell (RG) cells, which are located in the ventricular zone (VZ), constitute the major subtype of neurogenic cortical progenitors (Fietz and Huttner, 2010; Gotz and Huttner, 2005; Kriegstein and Alvarez-Buylla, 2009; Pinto and Gotz, 2007). They undergo multiple rounds of asymmetric cell divisions, thereby enabling the generation of diverse types of neurons while maintaining a pool of progenitors, (Miyata et al., 2004; Noctor et al., 2004). In humans, RG cells are thought to go through an increased number of such cycles when compared with non human-primate or mouse (Geschwind and Rakic, 2013; Lukaszewicz et al., 2005; Otani et al., 2016). The capacity to generate more neurons through a prolonged period is likely linked to species-specific properties intrinsic to the RG cells, but overall the underlying molecular mechanisms remain largely unknown. Observed differences in cortical neurogenic output between species are also linked to the expansion of specific classes of progenitors in the primate/human cortex, in particular the “outer” radial glial (oRG) cells, located in the outer-subventricular zone (OSVZ) (Fietz et al., 2010a; Hansen et al., 2010b; Reillo et al., 2011). oRG cells emerge from RG cells later in embryogenesis and their progeny tend to undergo multiple rounds of divisions, thus providing an additional mechanism of increased neuronal output.

From a molecular viewpoint, many highly conserved signalling pathways are required for the control of cortical progenitors amplification, differentiation, and survival, as well as neuronal specification (Gotz and Sommer, 2005; Molyneaux et al., 2007; Tiberi et al., 2012b). Interestingly, several differences have been identified between mouse and human and have been proposed to participate to species-specific differences in cortical neurogenesis including regulation of the Wnt (Boyd et al., 2015), SHH (Wang et al., 2016), PDGF (Lui et al., 2014), and Notch pathways (Rani et al., 2016).

The advent of detailed comparative analyses of mammalian genomes and transcriptomes, in particular those of hominids and primates, has enabled the identification of many human-specific candidate signatures of divergence, which might underlie some aspects of human brain evolution (Enard, 2016; Hill and Walsh, 2005; O’Bleness et al., 2012; Varki et al., 2008). One major driver of phenotypic evolution relates to changes in the mechanisms controlling gene expression (Carroll, 2003). Indeed, transcriptome analyses have started to reveal divergent gene expression patterns in the developing human brain compared with non-human species (Bae et al., 2015; Hawrylycz et al., 2012; Johnson et al., 2009; Johnson et al., 2015; Khaitovich et al., 2006; Lambert et al., 2011; Miller et al., 2014; Mora-Bermudez et al., 2016; Nord et al., 2015; Reilly and Noonan, 2016; Sun et al., 2005). Studies focused on the evolution of non-coding regulatory elements have further revealed structural changes that could lead to species-specific patterns of gene expression in the developing human brain (Ataman et al., 2016; Boyd et al., 2015; Doan et al., 2016; Pollard et al., 2006; Prabhakar et al., 2006; Prabhakar et al., 2008; Reilly et al., 2015). Changes at the level of coding sequences have also been proposed to contribute to human brain evolution, most strikingly the FoxP2 gene (Enard et al., 2002), encoding a transcription factor required for speech production that has acquired novel molecular targets (Konopka et al., 2009).

Another important driver of evolution is the emergence of novel genes (Ohno, 1999). Gene duplication, whether DNA or RNA-mediated (Kaessmann, 2010), is one of the primary forces by which novel gene function can arise, where an “ancestral” gene is duplicated one or several times into several related “paralog” genes (Dennis and Eichler, 2016; O’Bleness et al., 2012; Varki et al., 2008). Among these, hominid-specific duplicated (HS) genes, which arose from segmental DNA-mediated gene duplications specifically in the hominid and/or human genomes, are particularly interesting (Fortna et al., 2004; Goidts et al., 2006; Marques-Bonet et al., 2009; Sudmant et al., 2010). Indeed, most of them have occurred recently and specifically in the human/hominid lineage after its separation from the common ancestor to great apes such as Chimpanzees and Bonobos, during the period of rapid expansion of the cerebral cortex. They could inherently lead to considerable gene diversification and modification (through truncations, fusions, rapid sequence divergence, or novel transcriptional regulation of the duplicated paralog) and thereby may have contributed to the rapid emergence of human-specific neural traits. The role of the vast majority of the genes found in large segmental duplications remains unknown, and many of them could be non-functional, redundant with their ancestral form, or involved in non-neural aspects of human biology. Interestingly, recent segmental duplications are enriched for gene families with potential roles in neural development (Fortna et al., 2004; Sudmant et al., 2010; Zhang et al., 2011). Moreover, most human-specific gene duplications are found in recombination ‘hotspots’ displaying increased susceptibility to copy number variation (CNV) often linked to neurodevelopmental disorders, including changes in brain size (micro/macrocephaly), and/or altered cognitive development (mental retardation and autism-related syndromes) (Coe et al., 2012; Mefford and Eichler, 2009; Nuttle et al., 2016; Varki et al., 2008). Finally, recent studies have started to provide more direct evidence for the functional importance of HS gene duplications, including SRGAP2, ARHGAP11 and TBC1D3 (Charrier et al., 2012; Florio et al., 2015; Ju et al., 2016). These studies provide the first examples of HS gene duplications that may be linked to human cortex evolution, but it remains unclear how many and which HS genes are actually involved in corticogenesis. One of the roadblocks in identifying candidate HS genes that might play significant roles in cortical neurogenesis is that it remains very challenging to distinguish the expression of mRNA expressed from the ancestral gene or the HS paralogs, since their degree of conservation at the base pair level within exonic sequences is usually extremely high (Sudmant et al., 2010).

To tackle this question systematically, we first sought to determine which part of the HS gene repertoire is actually expressed during human corticogenesis, using tailored RNAseq analysis aimed at detection of HS gene expression with high specificity and sensitivity. We thus identified a specific repertoire of HS duplicated genes that show robust and dynamic patterns of expression during human fetal corticogenesis. In particular, we identified NOTCH2NL HS paralogs of the Notch receptor for their ability to promote cortical progenitor maintenance. Functional analyses of mouse and human cortical cells revealed that NOTCH2NL paralogs significantly expand human cortical progenitors and increase their neuronal output at the clonal level, through activation of the Notch pathway, mediated by inhibition of Delta/Notch cis interactions. Our results identify a novel cellular and molecular mechanism acting as a human-specific modifier of cortical neurogenesis through modulation of NOTCH signaling.

## RESULTS

### Identification of a repertoire of hominid-specific gene duplications dynamically expressed during human corticogenesis

Previous work has identified dozens of HS gene families containing genes duplicated in the hominid/human lineage (Dennis et al., 2017; Dumas et al., 2007; Fortna et al., 2004; Sudmant et al., 2010), but very little information is available on expression pattern in the developing human brain. This is largely due to the difficulty in assessing their expression with conventional methods, given the high sequence similarity between HS paralogs of the same family.

We first sought to determine, for each HS gene family, whether and how ancestral and paralog genes are actually expressed in the human developing cortex. We performed deep sequencing of RNA (RNAseq) extracted from human fetal cortex, at stages encompassing cortical neurogenesis (from 7 to 21 gestational weeks (GW)). For the later stage samples we also performed microdissection to disciminate specific regions of the cortex (frontal to occipital). In addition for the parietal area we microdissected the cortical plate (CP) and underlying domains of the cortical wall (non-CP, containing mostly OSVZ and VZ germinal zones)) in order to isolate compartments enriched in neural progenitors vs. postmitotic neurons.

To maximize the sensitivity and specificity of HS genes detection, we selected the libraries for cDNA fragments (>~350bp) and sequenced them with a 2×151bp paired-ends protocol (Figure S1A). Since duplications of HS genes are recent evolutionary events, paralogs within each HS gene families are highly similar, potentially confusing the mapping of reads originating from individual paralogs and subsequent estimates of their relative levels of expression (Figure S1B). Standard annotations on reference genomes (GRCh38/hg38) are also discordant for HS genes. We therefore manually curated gene structures for these genes whenever possible (Table S1) and developed a computational pipeline correcting the expression estimates of closely related paralogs for mapping errors (Figure S1C,D) (STAR Methods).

The distribution of gene expression values for HS genes was similar to that of all genes in the human reference genome (Figure 1A). We then selected HS genes based on their absolute levels of expression above a defined threshold (5 FPKM) corresponding to the minimal level of a set of 60 well-defined marker genes of cortical identity (Figure 1A). In addition, we selected the gene families where at least two paralogs, typically the ancestor plus at least one HS paralog, were expressed higher than such level (Figure 1B).

**Figure 1.**
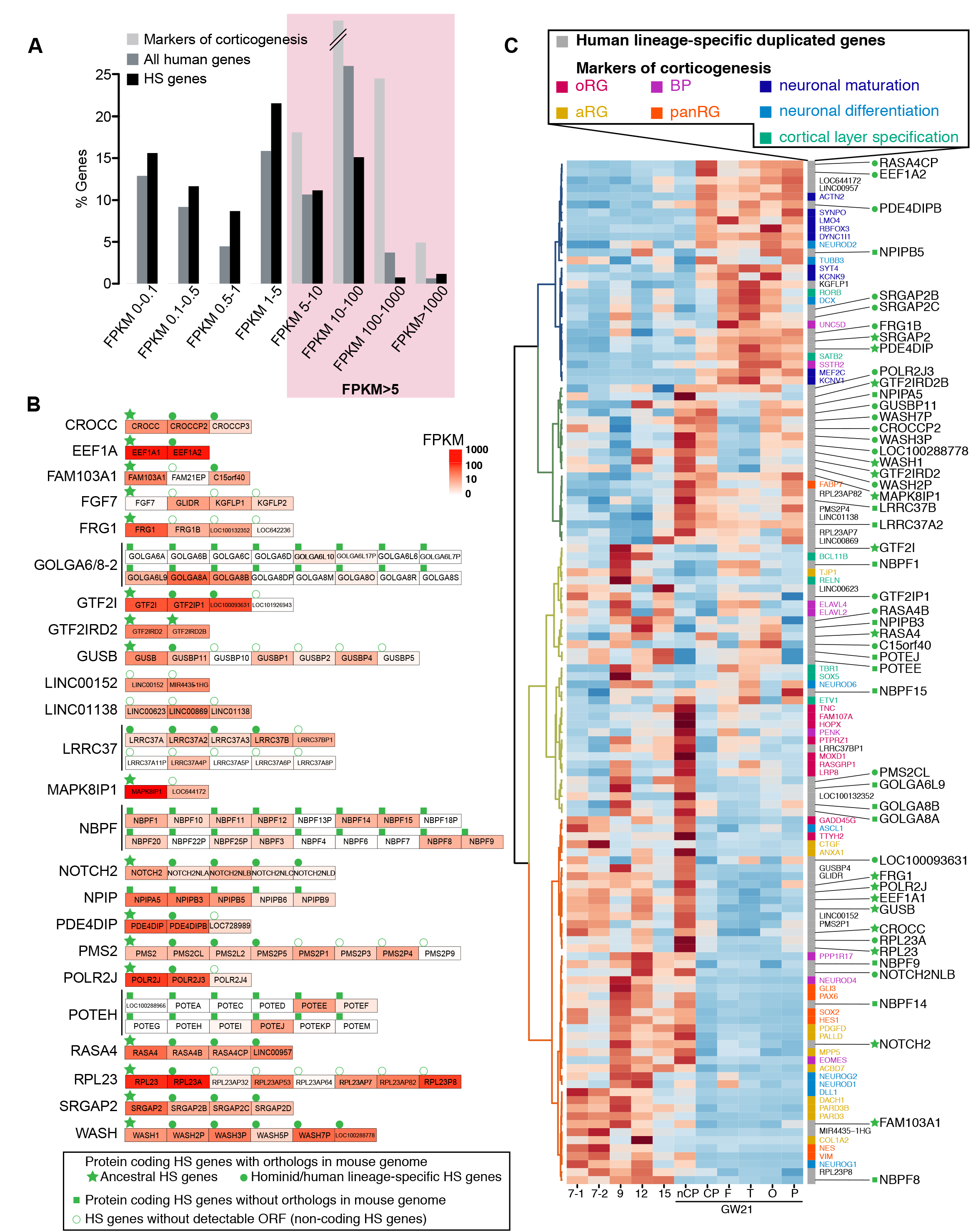
Transcriptome screening of HS genes expressed significantly during human corticogenesis. (A) Distribution of expression levels of HS genes is similar to that of all human genes among samples of fetal cortex. 60 selected marker genes of corticogenesis all display an expression level higher than FPKM5. (B) 24 families of HS genes identified by the transcriptome screen; color intensities represent the peak expression observed during the course of human fetal corticogenesis. For HS gene families where orthologue is present in the mouse, the ancestor is tagged with a green star, while HS paralogs are tagged with a green circle. HS genes without any ortholog in mouse are tagged with a green square. HS genes with no detectable ORF are tagged with a white circle. (C) Expression pattern of HS genes in the human fetal transcriptome; cluster analysis of HS and cortical marker gene expression in human fetal cortex. Samples are displayed horizontally from early to late stages in gestational weeks (GW) – at GW21 specific subdomains were isolated: non-cortical plate (non CP), cortical plate (CP), frontal (F), temporal (T), occipital (O) and parietal (P). Different colors identify HS genes (gray) and marker genes, such as aRG (apical radial glia), oRG (outer radial glia), panRG (all radial glia) and BP (basal progenitor). Corresponding color fonts for cortical marker genes are displayed in the left hand column. HS genes without an ORF are displayed in gray/blackand coding HS family genes are displayed in the right column. Heatmap colors were scaled for each individual row to better highlight temporal dynamics. The scale of expression for each HS gene corresponds to panel B.

This selection led to a short-list of 68 genes (including 17 potential ancestral genes) distributed among 24 gene families, displaying robust and temporally dynamic expression during human corticogenesis (Figure 1B and Table S2). HS genes displayed diverse patterns of gene expression, with about half of the genes preferentially expressed in progenitors/early stages, and fewer of them expressed preferentially in neuronal compartment/late stages of cortical neurogenesis (Figure 1C). Analysis of predicted coding sequences revealed that 51 HS genes (including 17 potential ancestor genes and 36 potentially unique to hominid/human genomes) display an open reading frame that was overall conserved, though distinct, between paralogs of the same family, strongly suggesting that they encode related but distinct proteins. For 17 HS genes, no conserved ORF could be detected, indicating potential pseudogenization or function as non-coding RNA (Table S2). Among the HS genes that have been validated so far in the context of cortical development, our screening successfully identified SRGAP2 family genes, but not ARHGAP11 and TBC1D3 families, because of the detectable expression of ARHGAP11B but below our threshold, and unreliable annotation for TBC1D3 family genes.

The cell diversity and spatial specificity of expression patterns was further tested by in situ hybridization (ISH) in human fetal cortex samples encompassing stages from 9 to 17 GW for 21/24 families (Figure 2). Specific ISH probes for paralogs were used whenever possible but in most cases probes recognizing more than one paralog had to be used. Significant expression in human fetal cortex was confirmed for 19/21 families (Figure 2 and data not shown), and revealed selective patterns of expression at the tissue and cellular levels.

**Figure 2.**
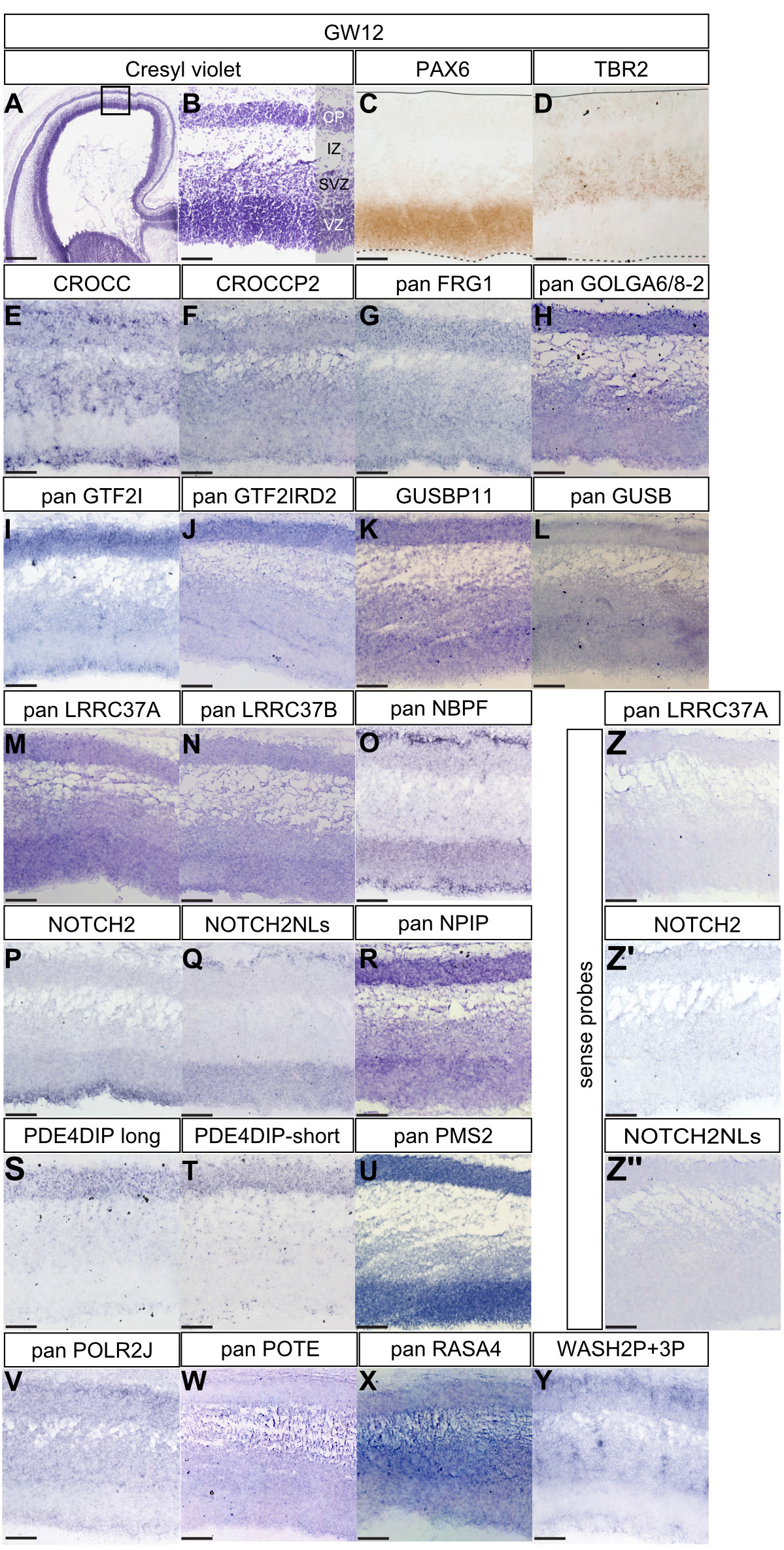
*In situ* hybridazation analysis of HS gene expression in the human fetal cortex. (A, B) Cresyl violet staining of coronal sections of gestational week (GW) 12 human fetal cortex. The dorsal region of the cortex (box in A) is enlarged in B. Adjacent sections are immunostained for apical progenitor marker PAX6 in the ventricular zone (VZ) and the basal progenitor marker TBR2 in the subventricular zone (SVZ) respectively (C, D). (E-Y) *in situ* hybridization of 16 selected HS gene families in GW12 human fetal cortex. No detectable signal is found when using sense probes (Z). Scale bars; 500μm (A) and 100μm (B).

Our data thus identify a specific repertoire of 36 protein-encoding genes that are potentially unique to hominid/human genome and are dynamically expressed during human corticogenesis. Their patterns of expression suggest potential involvement in multiple processes, from the control of progenitor biology and neurogenesis, to the regulation of neuronal migration, neuronal maturation and circuit formation.

### NOTCH2NL are four NOTCH2 paralogs expressed during corticogenesis

Based on these expression data, we next focused on HS genes expressed in cortical progenitors at the time of active neurogenesis, i.e. HS paralogs of the CROCC, GOLGA6/8, LRRC37A/B, NBPF, NOTCH2NL, NPIP, PDE4DIP, PMS2 and WASH gene families (Table S1). For each HS paralog of interest, we performed a small scale in vivo gain of function screen in the embryonic mouse cortex using in utero electroporation, looking at the distribution of electroporated cells between the ventricular zone (VZ) containing progenitors and cortical plate (CP) containing differentiated neurons. While changes in cell distribution and/or morphology outside the VZ were noted for some of the genes (data not shown), one particular family member, NOTCH2 N terminal like B (NOTCH2NLB), a Notch2 paralog, stood out because of its significant ability to maintain cells in the VZ compartment, thus potentially in the progenitor state (Figure 3A-D). NOTCH2NL was thus studied further to test whether and how it could function during human corticogenesis, initially focusing on gene structure and products of NOTCH2 HS paralogs.

**Figure 3.**
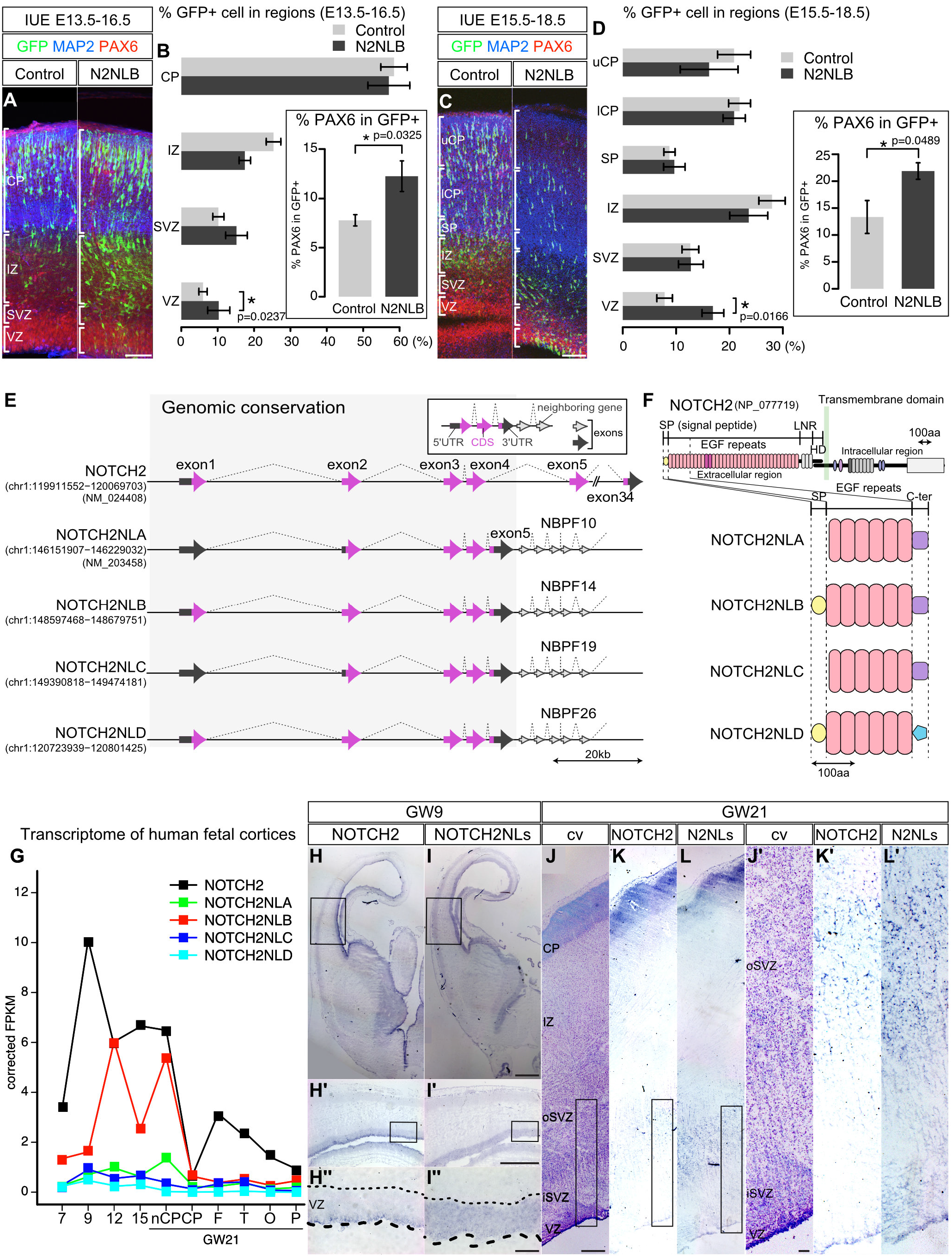
NOTCH2NL identification, structure and expression during corticogenesis. (A, B) *In utero* electroporation of NOTCH2NLB (N2NLB) in E13.5 and E15.5 mouse cortex. 3 days after electroporation, the fraction of electroporated cells in four regions(CP, IZ, SVZ and VZ), were collated. The percentage of PAX6-expressing cells among all electroporated cells was then calculated (B, D). Error bars = mean±sem, p values = Student’s t-test. (E) Structure of NOTCH2 gene and its four HS paralogs. Protein coding and non-protein coding exons are depicted as arrows in magenta and gray, respectively. (F) Predicted protein structure of NOTCH2NL members. SP (yellow) secretion signal peptide; EGF repeats (pink) (G) RNAseq expression pattern of NOTCH2-family genes during human corticogenesis. (H-L) *in situ* hybridization using NOTCH2-specific and NOTCH2NL-specific probes at 9 (H and I) and 21 (K and L) gestational weeks (GW). VZ ventricular zone, oSVZ outer subventricular zone, IZ intermediate zone, CP cortical plate. Scale bars; 100μm (A, C, I’’, J and J’), 500μm (I’’) and 1mm (I).

Detailed inspection of the reference genome (GRCh38/hg38) revealed that in addition to NOTCH2, which is present in all sequenced mammalian genomes, we found four additional NOTCH2 paralogs in the human genome, in two loci of chromosome 1 (Figure S2A). Among four putative NOTCH2NL loci, only one has a gene annotation in GRCh38/hg38, which is based on the transcript (NM_203458) identified in a previous study (Duan et al., 2004). We named the annotated NOTCH2NL locus as NOTCH2NLA, and named NOTCH2NLB, C and D the other three paralogs (Figure S2A and Figure 3E). Analysis of the genomic regions revealed that all four NOTCH2NL paralogs corresponded to the first 4 exons and introns of NOTCH2, thus the likely result of segmental duplications of the NOTCH2 ancestral gene (Figure 3E). In addition they contained a fifth exon unique to NOTCH2NLs, corresponding to an intronic region in NOTCH2. Interestingly three of the NOTCH2NL paralogs (A-C), as well as several other HS genes identified here above as expressed during corticogenesis (including NBPF11, NBPF12, NBPF14 and LINC01138), are found within the 1q21.1 region (Figure S2A), very close to a locus presenting CNV associated with microcephaly and macrocephaly (Brunetti-Pierri et al., 2008; Diskin et al., 2009; Dougherty et al., 2017; Mefford et al., 2008; Stefansson et al., 2014; Stefansson et al., 2008; Stone et al., 2008).

All NOTCH2NL paralogs are predicted to contain an open reading frame homologous to NOTCH2, but truncated and corresponding to the N terminal region of NOTCH2 extracellular domain (Figure 3F and Figure S2B-C). Each of the four encoded NOTCH2NLA-D proteins encode the 6 first EGF repeats of NOTCH2, followed by a C terminus domain not found in ancestral NOTCH2, and preceded for two of them by a predicted signal peptide. Inspection of publicly available genome sequence of other hominids and mammals revealed that 3/4 NOTCH2NL were detected exclusively in the human genome, while (at least) one was detectable in the chimpanzee genome, suggesting that NOTCH2NL duplications emerged very recently during hominid evolution.

RNAseq analyses of NOTCH2 and its paralogs revealed similar but distinct patterns of expression (Figure 3G). NOTCH2 was highly expressed at all stages of corticogenesis examined, with a peak at 9GW. NOTCH2NLB was expressed at lower levels at 7-9GW and then its levels increased at later stages, including in the non-CP region at GW21, containing the OSVZ. A similar trend was observed for NOTCH2NLA, although its levels were overall lower. NOTCH2NLC and D displayed detectable but very low (<1FPKM) levels of expression throughout corticogenesis (Figure 3G). The relative amount of expression of each paralog (B>A>>C>D) was also confirmed with qRTPCR with Notch2NL-specific primers followed by direct sequencing (data not shown). Expression of NOTCH2 and NOTCH2NL paralogs was examined in more detail by *in situ* hybridization, using a probe recognizing potentially all four NOTCH2NL paralogs (and not the ancestral NOTCH2), and another probe specific to the ancestral NOTCH2 mRNA (Figure 3H-L). This revealed similar but distinct patterns of expression in VZ at early stages (GW9), with NOTCH2 expressed mostly along the apical part of the VZ, while NOTCH2NL genes were expressed throughout the VZ, in a salt and pepper pattern at GW9 (Figure 3H and I) and GW12 (Figure 2P-Q). At later stages (GW21) both NOTCH2 and NOTCH2NL genes appeared to be expressed heavily in a salt and pepper pattern, mostly in the outer subventricular zone that contains oRG cells (Figure 3J-L). These data indicate that NOTCH2 and its paralogs appear to be expressed in similar ways throughout corticogenesis, both in VZ (containing RG cells) and OSVZ (containing oRG cells).

### NOTCH2NLB leads to clonal expansion of human cortical progenitors

Given the prominent role of Notch signaling in controling the balance between selfrenewal and differentiation of neural progenitors throughout the animal kingdom (Kageyama et al., 2009; Kopan and Ilagan, 2009), we hypothesized that NOTCH2NL genes could represents hominid-specific modifier of cortical neurogenesis. Given the prominent expression of NOTCH2NLB and its impact detected by *in utero* electroporation on mouse cortical progenitors, we focused on this specific paralog for more detailed analysis.

In order to characterize the role of NOTCH2NLB in human cortical progenitors in a highly physiologically relevant manner, we used an in vitro model of cortical neurogenesis from human embryonic stem cells (Espuny-camacho et al., 2013). Importantly, in this system the species-specific temporal dynamics of cortical neurogenesis observed in vivo is recapitulated in vitro, with human and mouse cortical neurogenesis taking place over about 12 and 1 week, respectively. In the human system, NOTCH2 was expressed at high levels from the onset of in vitro corticogenesis, while NOTCH2NL gene expression steadily increased until two months of differentiation, thus reminiscent of the in vivo pattern of expression described above (data not shown).

We first tested the impact of NOTCH2NLB on cortical neurogenesis using a gain of function approach, focusing on the early stages where it is not expressed at highest levels yet. To achieve maximal sensitivity and specificity, we used a lentiviral-based clonal analysis assay allowing to measure the potential effect of NOTCH2NL on clonal amplification and differentiation from single labeled cortical progenitors (Figure 4A and B).

**Figure 4.**
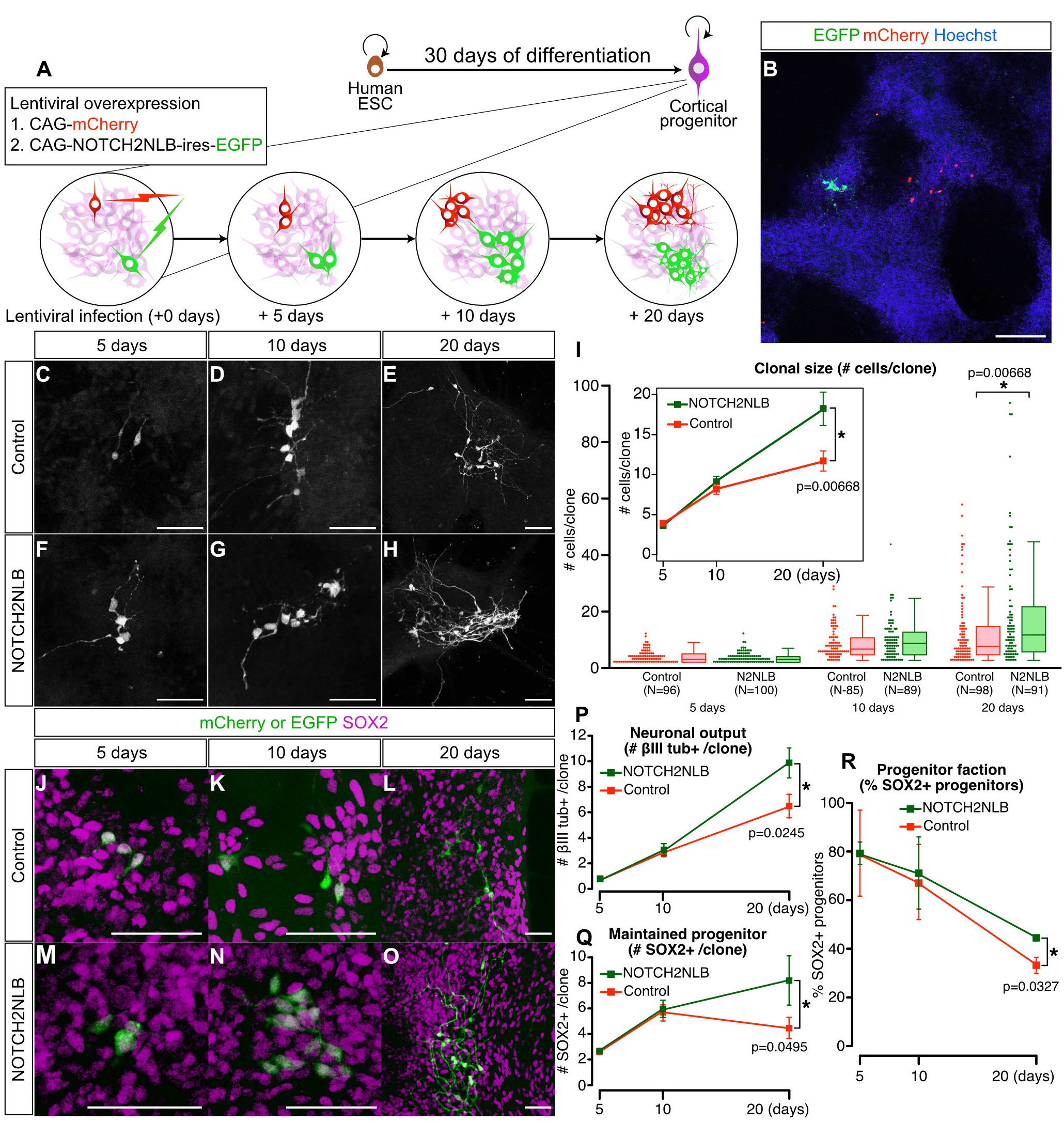
NOTCH2NLB overexpression leads to clonal expansion of human cortical progenitors. (A, B) Schematic illustration of the clonal analysis. Human ESC are first differentiated into cortical cells until 30 days *in vitro*, followed by lentivirus infection allowing clonal analysis for up to three weeks. (B) Representative clones (10 days post infection). (C-I) Representative cases of control (C-E) or NOTCH2NLB (F-H) clones. (I) Quantification of clonal size (number of cells per clone) in control and folowing NOTCH2NLB-overexpression. Dotplots and boxplots indicate the number of cells per clone. (N = number of clones analyzed) Mean±sem of clonal size is indicated in the inset (topleft). (J-O) Expression of cortical progenitor marker SOX2 in control mCherry- and NOTCH2NLB clones. (P-R) The numbers of neuron and progenitors were quantified following immunostaining of βIII tubulin and SOX2, respectively. Error bars = mean±sem and p value by Student’s t-test. Scale bars; 100μm (B) and 50μm (C-H and J-O).

In control conditions, early cortical progenitors amplified gradually to lead to a doubling of the clone size after 20 days culture (Figure 4C-I), while during the same period the number of Sox2-positive progenitors per clone gradually diminished (Figure 4J-R). Remarkably, NOTCH2NLB expressing clones almost tripled in size during the same period, and the proportion of Sox2 progenitors kept increasing, thus reflecting an increased capacity of NOTCH2NLB-expressing progenitors to expand clonally (Figure 4M-R). Moreover, quantification of the number of neurons in each clone revealed that NOTCH2NLB clones led to a larger neuronal output than in controls (Figure 4P). These data indicate that NOTCH2NLB expression in cortical progenitors leads to larger clone size, slower exhaustion of progenitor pool, and ultimately a higher number of neurons generated. This strikingly parallels the expected features of human corticogenesis compared with non-human primates (Geschwind and Rakic, 2013; Lui et al., 2011; Molnar et al., 2006).

### NOTCH2NLB promotes progenitor cell cycle reentry, mainly through its EGF repeats

The effects of NOTCH2NLB on clonal amplification could, in principle, be due to selfrenewal/decreased differentiation, or increased proliferation. To distinguish between these possibilities, we examined the effect of NOTCH2NLB on cell fate and cell cycle properties following 7 days of NOTCH2NL overexpression (Figure 5A-F). This analysis revealed an acute effect of NOTCH2NLB leading to increase of progenitor fate (PAX6 expression) at the expense of neuronal fate (βIII tubulin expression). Moreover, mitotic index assessed by short term (1h) DNA labeling pulses by a nucleotide analog EdU, as well as M phase marker stainings, revealed no difference following NOTCH2NLB overexpression, suggesting no overt effect on mitotic rate (Figure S3A-E). However, cumulative (24h) EdU labeling combined with labeling for all cycling cells (Ki67) revealed a signficant decrease in cell cycle exit, and conversely an increase in cell cycle re-entry following expression of NOTCH2NLB (Figure S3F-I). These data indicate that NOTCH2NLB increases clonal expansion mainly through increased self-renewal but without overt effect on proliferative rate *per se*.

**Figure 5.**
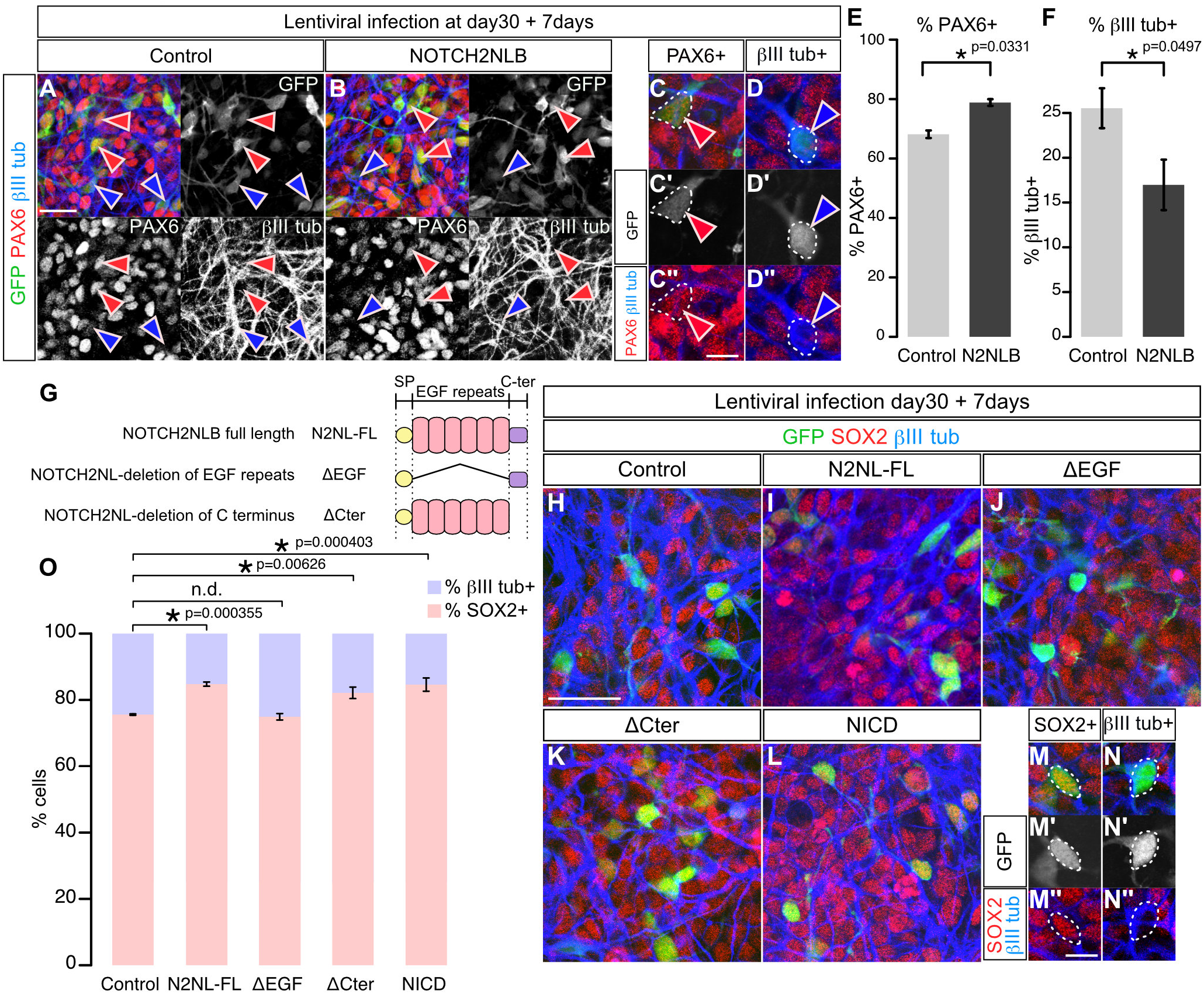
Overexpression of NOTCH2NLB in human cortical progenitors enhances progenitor maintenance through its EGF repeats. (A-F). Expression of NOTCH2NLB (N2NLB) (GFP+) in cortical cells 7 days post lenti-viral infection; the percentage of PAX6-expressing progenitor and βIII tubulin-expressing neurons was counted and analysed (E, F). (G) Predicted protein structures of the deletion mutants. (H-N) Representative images of SOX2 and βIII tubulin expression followinf infection with different constructs. (O) Quantification of the effects of the deletion constructs and NICD. Data are represented as mean±sem and p value by one-way ANOVA and bonferroni post hoc test. Scale bars; 50μm (A and H) and 10μm (C, D, M and N). Percentage of SOX2-expressing progenitor and βIII tubulin-expressing neuron are examined 7 days after lentiviral infection at day 30 of cortical differentiation of human ESC (O-N).

We next examined the molecular mechanisms underlying NOTCH2NLB function, by directly comparing the effect of NOTCH2NL mutants devoid of specific domains of the protein (Figure 5G-N). This revealed that the EGF repeats of NOTCH2NL are absolutely critical for their function, while the C-terminal domain appears to be dispensable. This was confirmed in the mouse embryonic cortex *in vivo*, where sustained cortical progenitor maintenance was found to be dependent on NOTCH2NL EGF repeats (Figure S3J-X).

### NOTCH2NLB directly promotes activation of the Notch pathway

As Notch signalling is known to lead to cortical progenitor self-renewal and block differentiation in the mouse and human cortical progenitors, we reasoned that the effects of NOTCH2NL could be linked to Notch activation (Kageyama et al., 2009; Lui et al., 2011). We tested this specifically by examining the effect of the canonical Notch pathway in the same in vitro paradigm on human cortical progenitors, using overexpression of Notch intracellular domain (NICD) that acts as an activator of the Notch pathway. This led to a similar effect as NOTCH2NLB, where progenitors numbers increased and the number of differentiated neurons decreased (Figure 5O).

These data suggest that NOTCH2NLs act through Notch pathway activation, in a way that is dependent on its EGF repeats. To test the impact of NOTCH2NLB on Notch signalling directly, we first examined the expression of Hes1, a direct downstream effector of the Notch pathway that is well-known to promote cortical progenitor self-renewal(Kageyama et al., 2008), in response to NOTCH2NLB and NICD expression (Figure S4). This revealed that acute expression of NOTCH2NLB, similarly to NICD, leads to upregulation of Hes1, and that this effect is dependant upon the presence of EGF repeats.

We then tested more directly for an interaction of NOTCH2NLB with the Notch pathway in vivo, by evaluating its effect on a classical Notch activity transcriptional reporter (CBFRE-EFGP) in cortical progenitors in vivo (Mizutani et al., 2007) (Figure 6A). This revealed that NOTCH2NLB overexpression in mouse cortical progenitors increases Notch reporter activity (Figure 6C and G), with an amplitude comparable to that induced by NICD (Figure 6C and I), while no effect was obtained with NOTCH2NL mutants devoid of EGF repeats (Figure 6C and H).

**Figure 6.**
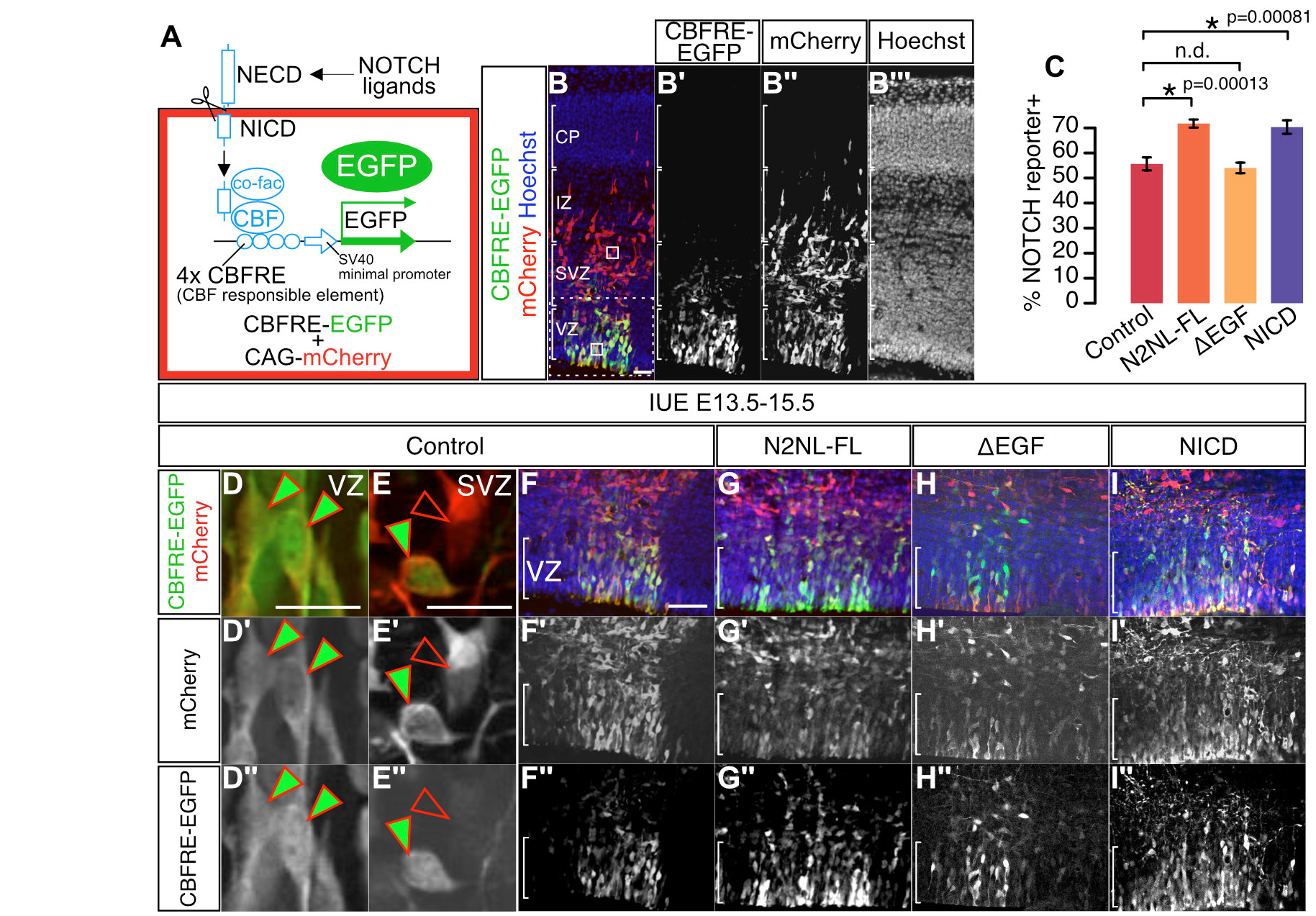
Overexpression of NOTCH2NLB upregulates NOTCH signaling *in vivo*. NOTCH signaling strength was investigated using a NOTCH reporter construct (CBFRE-EGFP) co-electroporated with control or NOTCH2NL overexpression constructs. (A) A NOTCH reporter containing CBF responsible element (CBFRE) drives expression of EGFP. (B) Cells in which Notch is activated can be identified as GFP+/mCherry+, while those where Notch is inactive are only mCherry+. (C-I). (C) *In utero* electroporation of NOTCH2NLB full length (N2NL-FL) and NICD increase NOTCH activation compared with control or NOTCH2NL deletion mutant of EGF repeats (ΔEGF). Data are represented as mean±sem and p value by one-way ANOVA and bonferroni post hoc test. Scale bars; 100μm (A and F) and 20μm (D and E).

### NOTCH2NLB acts through cell-autonomous inhibition of Delta/Notch interactions

Overall our data indicated that NOTCH2NLB appears to act mainly through activation of the Notch pathway, raising the question of the molecular mechanism involved. Since the EGF repeat domains are thought to bind to Notch ligands we first tested whether NOTCH2NLB could interact with canonical Notch ligand Delta-like 1 (DLL1), which is the major Notch ligand previously involved in cortical neurogenesis (Kageyama et al., 2008; Nelson et al., 2013; Shimojo et al., 2008). We hypothesized that NOTCH2NLB could directly effect DLL1 function and/or trafficking. For this purpose we used an inducible system of DLL1 induction, in which expression of DLL1 is transcriptionally independent of Notch (LeBon et al., 2014; Sprinzak et al., 2010), and measured the amount of functional DLL1 present at the plasma membrane and available for binding to soluble Notch1 extracellular domain added to the cell medium (Figure 7A). In these conditions, the amount of DLL1 functionally available at the plasma membrane was directly proportional of the total amount of DLL1 present throughout the cell, as expected (LeBon et al., 2014) (Figure 7A and B). Remarkably, the cells expressing both NOTCH2NLB and DLL1 displayed a strong reduction of the amount of Notch binding at the plasma membrane compared to control (Figure 7A-C), thus revealing a reduction of functional DLL1 expression at the cell surface, while co-expression of the NOTCH2NLB mutant devoid of EGF repeats was without effect on DLL1 (Figure 7B and C).

**Figure 7.**
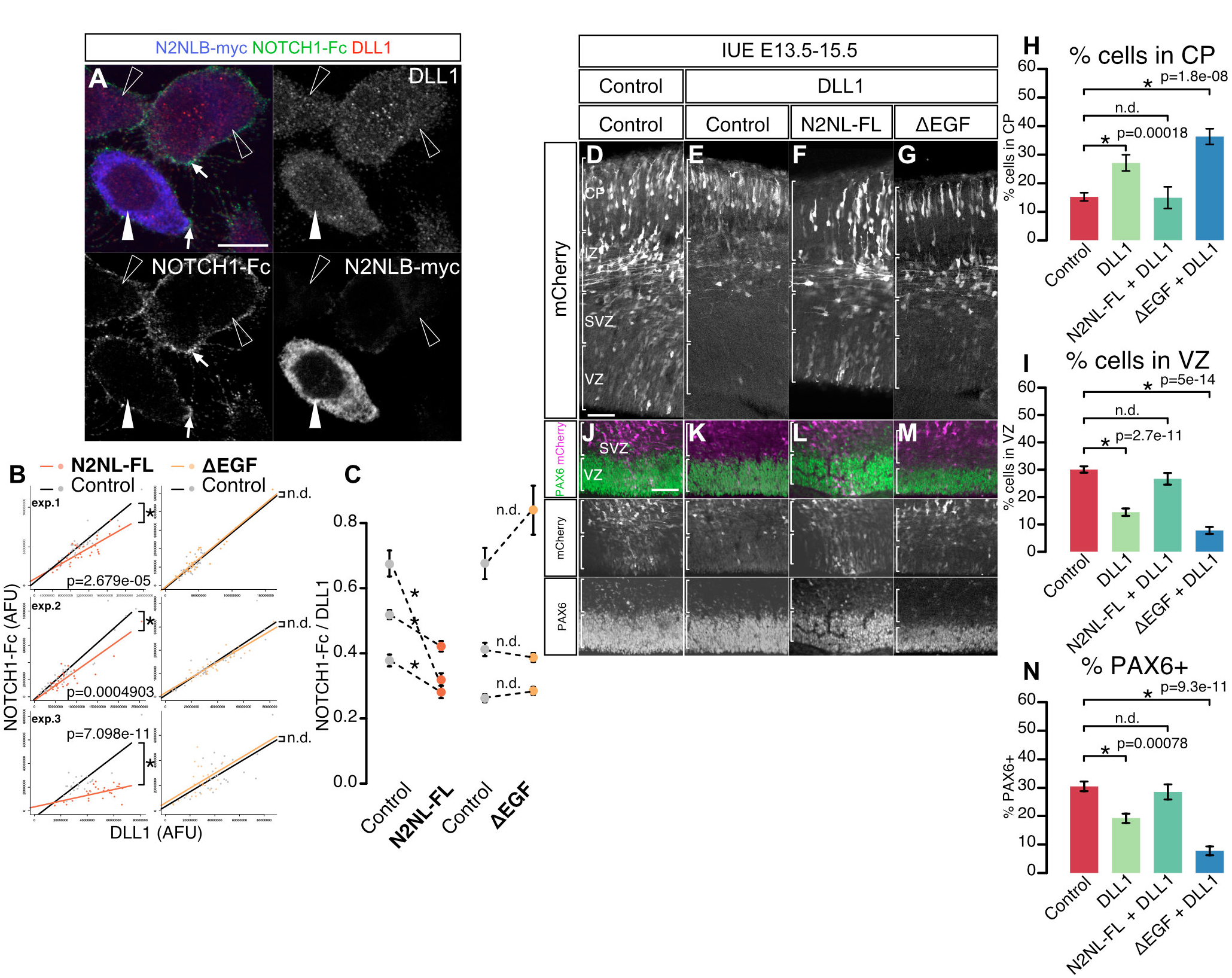
Cell-autonomous suppression of DLL1 function by NOTCH2NL. (A) A CHO cell line containing tetracycline-dependent expression of DLL1 was transfected with a NOTCH2NLB expression plasmid. DLL1 protein on the plasma membrane was detected by NOTCH1-Fc (green in A and white arrows)applied to living cells before fixation. Cells were immunostained using anti-DLL1 antibody to detect all DLL1 protein in the whole cell (red in A). Anti-Myc antibody was used to identify NOTCH2NLB-expressing cells (blue in A and white arrowhead) from the surrounding non-expressing cells (open arrowheads). (B and C). The fluorescent intensities of NOTCH1-Fc and of DLL1 are measured in both NOTCH2NLB-expressing cells and non-expressing cells. Plots of three independent experiments are shown. Compared to the non-expressing cells as negative control, NOTCH2NLB full length (N2NL-FL)-expressing cells show lower level of NOTCH-Fc signal, while NOTCH2NL-ΔEGF expressing cells display same levels as control cells (B). The ratio of signals for NOTCH1-Fc over DLL1 is quantified for each individual experiment (C). (D-N) N2NL-FL suppresses DLL1 function *in vivo*. Mouse *in utero* electroporation was performed with DLL1 alone, or together with N2NL-FL/ΔEGF. Bin analysis reveals that the proportion of electoporated cells is in increased in CP (H) and decreased in VZ (I) following DLL1 overexpression. This effect is abrogated by N2NL-FL co-expression, while ΔEGF has no effect (D-I). The same results are obtained when examining the proportion of PAX6 progenitors (J-N). Data are presented as mean ± SEM and p values by Student’s t-test (B and C) and one-way ANOVA and bonferroni post hoc test (H, I and N). Scale bars; 10μm (B) and 100μm (E, K).

These data indicate that NOTCH2NLB directly impacts, in a cell-autonomous fashion, the amount of DLL1 functionally available at the membrane. This, in turn, could lead to a reduction of DLL1 function in the NOTCH2NL-expressing cell. We tested this hypothesis and examined whether NOTCH2NL could lead to inhibition of DLL1 function during cortical neurogenesis. Indeed, DLL1 expression was previously shown to exert a cell-autonomous promotion of neuronal fate and decreased progenitor self-renewal in the mouse cortex in vivo (Kawaguchi et al., 2008). This suggests that NOTCH2NLB can promote progenitor self-renewal through cell-autonomous inhibition of DLL1 in cortical progenitors. We therefore examined the functional interactions between NOTCH2NLB and DLL1, by comparing the effect of DLL1 expression during mouse corticogenesis in vivo, alone or in combination with NOTCH2NLB (Figure 7D-N). As predicted (Kawaguchi et al., 2008), DLL1 overexpression led to a decreased proportion of progenitor cells in the VZ (Figure 7D, E, I, J and K), together and a corresponding increase in neurons in the post-mitotic compartments (CP; Figure 7D, E, H, J and K). This DLL1-mediated effect was completely blocked by NOTCH2NLB co-expression (Figure 7F, H, I, L and N), while the EGF repeat mutant could not counteract the DLL1 effects (Figure 7G, H, I, M and N). Overall our data demonstrate that NOTCH2NLB can directly inhibit DLL1 function cell-autonomously, leading to cortical progenitor expansion and self-renewal.

## Discussion

Gene duplication represents a driving force during evolution (Ohno, 1999), but it remains unclear how many HS duplicated genes may contribute to human brain evolution. The functional impact of HS gene duplication is also largely unknown, especially since the expression pattern of HS genes is very delicate to determine in an accurate way, given the strong homology between related paralogs and between paralogs and their potential ancestor genes.

Here, we used RNAseq tailored specifically to tackle this issue and detect HS duplicated genes with greater sensitivity and specificity. We thus determined which HS genes are expressed during corticogenesis, among which we identified a specific repertoire of 36 protein-encoding HS paralogs characterized by robust and dynamic expression patterns during human corticogenesis. Our data constitutes a rich resource to test the role of candidate HS modifiers of cortical neurogenesis, neuronal maturation and circuit formation. In addition some of the genes that are expressed below our fairly stringent threshold may be of functional interest, such as the recently identified ARHGAP11B (Florio et al., 2015). Nonetheless our analysis suggests that the vast majority of HS genes are expressed at low or undetectable levels in the human fetal cortex, which is in line with the predicted outcome of most gene duplications. This implies that most of these are unlikely to be involved in human corticogenesis, at least in a way that is intrinsic to the cortex. Besides, several of these HS paralogs do not appear to encode proteins, suggesting they might be pseudogenes or encode noncoding RNAs.

Among the HS genes identified in this study, we characterized functionally one important family of four paralogs of NOTCH2, NOTCH2NL, which act through the Notch pathway to increase self-renewal and ultimately neuronal output of human cortical progenitors. The recent evolutionary emergence of NOTCH2NL paralogs makes them attractive candidates to be involved in the latest aspects of human brain evolution, including increased size and complexity of the cerebral cortex. On this basis our data suggest that NOTCH2NL functional paralogs may have been selected as positive regulators of the well-conserved Notch pathway, and thereby could contribute in an important way to the evolution of human cortical neurogenesis, which is characterized by greater clonal expansion and higher rates of self-renewal of cortical progenitors (Dehay and Kennedy, 2007; Geschwind and Rakic, 2013; Lukaszewicz et al., 2005; Otani et al., 2016).

Functionally, we demonstrate that NOTCH2NLB has a critical functional impact on clonal expansion of human cortical progenitors, though a direct effect on their selfrenewal. This effect is strikingly in line with the expected pattern of human cortical neurogenesis compared with non-human primates: indeed, human cortical RG cells are expected to go through an increased number of self-renewing cycles when compared with non-human primate species (Dehay and Kennedy, 2007; Geschwind and Rakic, 2013; Lukaszewicz et al., 2005; Otani et al., 2016). Importantly, the capacity to generate neurons for a prolonged period is likely linked to species-specific properties intrinsic to the RG cells, since pluripotent stem cells-derived RG cells from mouse, macaque and human follow a species-specific temporal pattern of neurogenesis similar to in vivo (Espuny-camacho et al., 2013; Otani et al., 2016), and the increased clonal amplification and self-renewal potential of human vs. non-human primate RG cells is retained in vitro (Otani et al., 2016). Here we identify NOTCH2NL as species-specific regulators of such processes, acting by increasing the level of self-renewal of RG cells without detectable effect on cell cycle dynamics per se. Through these effects Notch2NL could thus enable a longer period of neurogenesis and larger neuronal output that is characteristic of human/hominin lineages.

Interestingly NOTCH2BNLA/B appear to be also expressed at high levels in oRG cells, which have been proposed to play an important role in the increase of cortical size in primates and other species (Fietz et al., 2010a; Hansen et al., 2010b; Reillo et al., 2011). Although we have not looked at the specific impact of NOTCH2NL on oRG cells (which appear much later during in vitro corticogenesis, making them harder to investigate using clonal analyses), our data on RG cells strongly suggest that Notch2NL could contribute to the expanded self-renewal capacity of oRG cells as well, inasmuch that their self-renewal was shown to depend on Notch signalling (Hansen et al. 2010). Noteworthy, we do not observe increased number of oRG cells or other types of basal progenitors in the mouse cortical progenitors following overexpression of NOTCH2NLB, implying that it mostly acts by increasing the self-renewal capacity of apical RG cells, and not by promoting the generation of basal progenitors. NOTCH2NL has thus a different function from the one proposed for HS genes ARHGAP11B and TBC1D3 that were shown to promote generation of basal progenitors (Florio et al., 2015; Ju et al., 2016).

At the molecular level, we found that the effects of NOTCH2NL are strikingly similar to the effects of Notch, both in vitro and in vivo, and indeed our data also indicate that NOTCH2NL appear to act mostly through the direct stimulation of Notch signalling. NOTCH2NL thus impinges on one of the most conserved developmental pathways controlling cell fate in embryonic development (Kageyama et al., 2009; Kopan and Ilagan, 2009). Given the paramount functional importance of the Notch pathway during neurogenesis, and its multiple levels of regulation (Bray and Bernard, 2010; Kageyama et al., 2009; Pierfelice et al., 2011), NOTCH2NL may be one of many species-specific regulators of the Notch pathway in the developing brain: indeed a recent study identified a primate-specific long non-coding RNA that appears to control positively the expression of Notch receptors (Rani et al., 2016).

Interestingly NOTCH2NL acts at least in part through the inhibition of Delta function, in a cell-autonomous fashion. Future work should determine the exact mechanism by which this leads to Notch activation: it could lead to Notch activation through trans-signalling from neighbouring cells, as predicted by lateral inhibition models, but it could equally act by direct blockade of the so-called Notch cis-inhibition process. Cis-inhibition is well described in several model systems (Bray, 1997; Cordle et al., 2008; del Álamo et al., 2011; Klein et al., 1997; LeBon et al., 2014; Matsuda and Chitnis, 2009; Micchelli et al., 1997; Miller et al., 2009; Sprinzak et al., 2010), where Delta acts as a direct inhibitor of the Notch receptor in the cells where they are co-expressed, thus in *cis*. As NOTCH2NLB appears to block Delta function when co-expressed in the same cell, a likely mechanism could be that it blocks Delta-mediated cis-inhibition, and thereby allows for a higher tone of Notch signalling, in a cell-autonomous manner. Future work should also enable to determine whether NOTCH2NLB impacts on Delta within the cell, thus regulating trafficking of Dll1, or at the level of the membrane, where it could interfere with Dll1 binding sites available to Notch in cis.

The pro-Notch effect of NOTCH2NL in Delta-expressing cells is particularly interesting in the proposed model of oscillations of the Notch pathway in cortical progenitors (Shimojo et al., 2008). Indeed in this model progenitors are thought to commit to neuronal fate only beyond a certain threshold of proneural gene expression and activity, and our data predict that this threshold would be set at a higher level in the presence of NOTCH2NL, given the blockade of Delta-mediated inhibition of Notch. According to this model, NOTCH2NL would act in Delta-expressing progenitors close to their final commitment to neuronal fate, and would contribute to keep them in an undifferentiated state.

This does not exclude however that NOTCH2NL may also act in a non-autonomous manner, through lateral inhibition or other mechanism, as it is predicted to be secreted, and it could also bind to other Notch ligands or receptors, through its EGF repeats that may mediate mediate both heterophilic and homophilic interactions.

In any case, it is interesting to note that the mechanism of action of NOTCH2NL, acting as a positive regulator of its ancestor NOTCH2, is quite distinct from the mechanisms proposed for SRGAP2 and ARHGAP11, for which Hs paralogs also encode a truncated form of the ancestral form. Indeed, for SRGAP2, the HS paralogs have been shown to act essentially as inhibitors of the ancestral form (Charrier et al., 2012), while ARHGAP11B is proposed to work independently of ARHGAP11A (Florio et al., 2015). Interestingly, many of the proteins encoded by the 35 HS genes identified in this study appear to encode truncated forms: it will thus be interesting to determine how they act in the context of corticogenesis and beyond.

Our data strongly suggest that NOTCH2NL paralogs could act as species-specific modifiers of cortical size. In line with this hypothesis, NOTCH2NLA/B/C paralogs are found within the 1q21.1 genomic region, within the interval of CNV associated with pathological changes of brain size and cognitive defects (Brunetti-Pierri et al., 2008; Diskin et al., 2009; Mefford et al., 2008; Stefansson et al., 2008; Stone et al., 2008). Most strikingly, microdeletions in this 1q21.1 locus can lead to microcephaly while microduplications can lead to macrocephaly: should NOTCH2NL paralogs be involved in these CNV their cellular effects would predict these effects on brain size, given the ability of NOTCH2NL to promote expansion of human cortical progenitors.

In striking convergence with our study in this context, a very recent study (Fiddes et al. *submitted*) has identified the same functional NOTCH2NL paralogs as resulting from very recent duplications in the hominin lineage, and precisely mapped NOTCH2NLA/B within the interval of several microdeletions and duplications in patients with the 1q21.1 CNV syndrome.

In conclusion, our study reveals a selective repertoire of hominid-specific gene duplications with a potentially high impact for human brain evolution, including NOTCH2NL paralogs that act as species-specific modifiers of Notch signalling and neurogenesis, with direct links with our recent genomic evolution, but also potentially with neurodevelopmental diseases that may strike specifically the uniquely large and complex human cerebral cortex.

## Acknowledgments

We thank members of the lab and IRIBHM for most helpful feedback and support, and Jean-Marie Vanderwinden of the Light Microscopy Facility (LiMiF) for support with imaging. We thank Drs. Lambot and Lambert and the clinics of obstetrics and gynecology of Erasme hospital for help with collection of fetal material. We are grateful to Dr. Evan Eichler for insightful advice and discussions. This work was funded by grants from the Belgian FNRS, the European Research Council (ERCAdv Grant Gendevocortex), the Belgian Queen Elizabeth Medical Foundation, the Interuniversity Attraction Poles Program (IUAP), the WELBIO Program of the Walloon Region, the Fondation de Spoelbergh, the AXA Research Fund, and Fondation ULB, and the ERA-net E-Rare ‘Microkin’ (to P.V.). I.K.S. and D.K. were supported by EMBO Fellowships. I.K.S. was a postdotoral Fellow of the FNRS, R.V. is an Aspirant of the FNRS and supported by a L’Oréal-Unesco For Women in Sciences fellowship, and M.W. is a PhD Fellow of the FRIA.

## Author contributions

I.K.S. performed all experiments, with the help of R.V., D.K., M.W., A.B. and A.H. D.G. and V.D. designed and performed computational analyses of RNASeq data. I.K.S. and P.V. designed and analyzed all experiments and wrote the manuscript, together with D.G., V.D., and F.P.

**STAR+METHODS OUTLINE**

**CONTACT FOR REAGENT AND RESOURCE SHARING**

**EXPERIMENTAL MODEL AND SUBJECT DETAILS**

**Human fetal tissue collection and preparation**
**Mice**

**METHOD DETAILS**

**RNA sequencing**
**Transcriptome analysis**
***In situ* hybridization**
**Digoxigenin (DIG)-labeled riboprobe preparation**
**ESC culture and differentiation**
**Lentiviral preparation**
**Clonal analyses**
**Acute overexpression in human cortical cells**
**Cell cycle labeling assay**
***In utero* electroporation**
**Constructs**
**Immunofluorescence staining**
**Detection of DLL1 protein on the plasma membrane**
**Confocal microscopy**

**QUANTIFICATION AND STATISTICAL ANALYSIS**

**Statistical analysis**
***In utero* electroporation in mouse**
**Clonal assay in human cortical cells**
**Detection of DLL1 protein on the plasma membrane**

**DATA AND SOFTWARE AVAILABILITY**

## Supplementary figures

**Figure S1 (Related to Figure 1).**
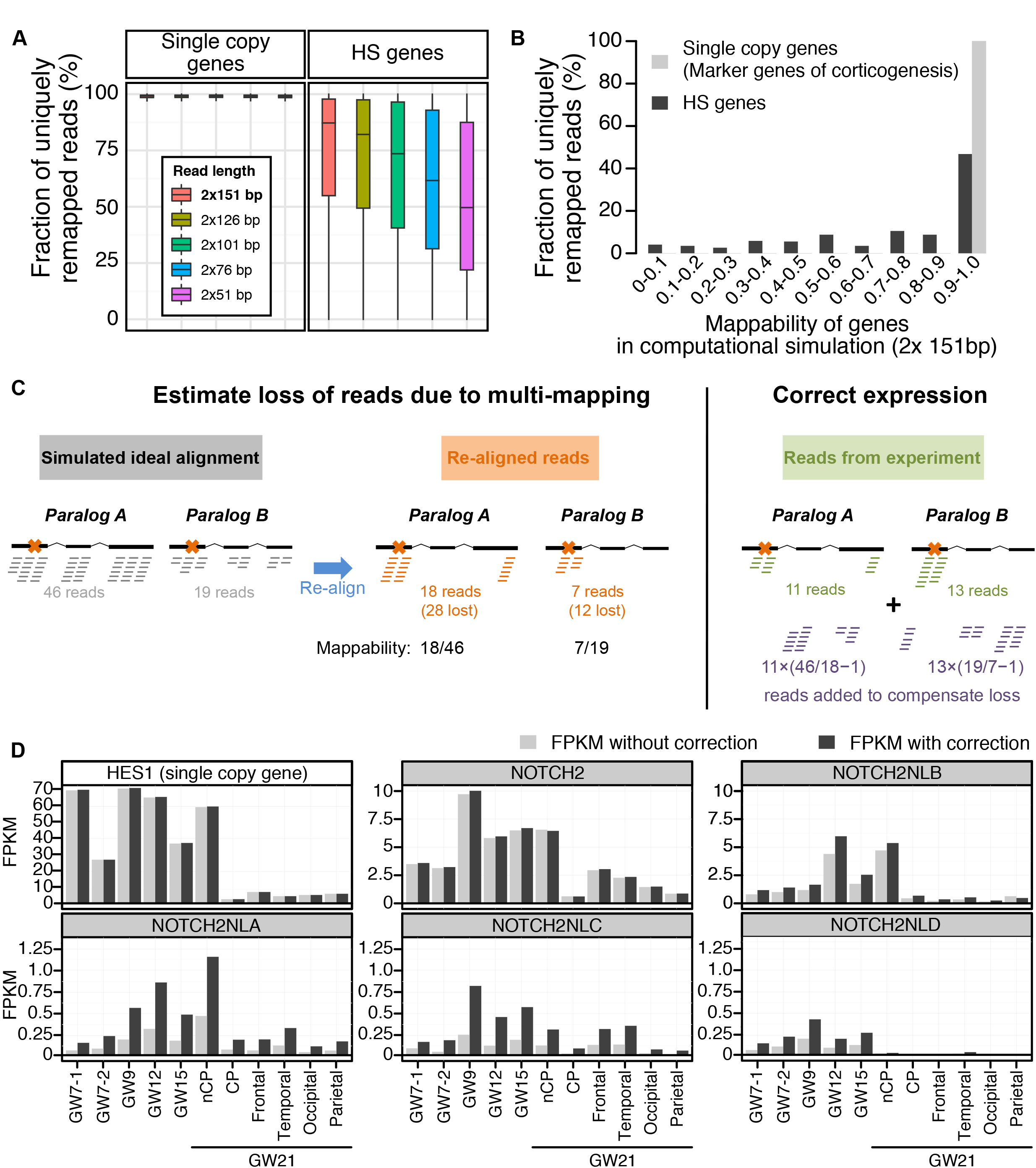
Transcriptome analysis: correcting for loss of multimapping loss. (A) The mappability of a gene is defined as the number of reads uniquely mapped on this gene divided by the number of reads originating from its transcripts. It is computed from simulated transcriptomes, as explained above and depicted in panel C. Most reads originating from single copy genes are uniquely mapped on the reference genome regardless of sequencing length, hence they have a mappability of one. HS genes show lower mappability values. As expected, their mappability is inversely related to read length. (B) When setting simulated read length to 2x 151bp, which is the length used for transcriptome sequencing of human fetal cortex, only half of the HS genes show mappability values above 90%. (C) Computational correction of expression for human specific paralogs. Paralogs within each HS gene families are highly similar; potentially confusing the mapping of reads originating from individual paralogs. As a result, some reads are discarded because they map to multiple paralogs, leading to expression under-estimation. To estimate this loss quantitatively, an alignment of simulated reads (BAM file) is generated for each gene (grey alignment) at a defined coverage (see methods). This simulated alignment is ideal as it assumes a uniform coverage of the genes, and importantly the reads are manufactured and placed on reference genome, i.e. no read mapping procedure is involved, hence there is no mapping ambiguity. These simulated reads are then extracted and aligned with the same alignment procedure as used for *in vivo* experimental data (see methods; orange alignment, crosses on the gene structures denote unique sequence features allowing unambiguous mapping). Many reads are lost in the process due to multimapping, but we can estimate how many, since we initially generated them in known quantity (i.e. grey alignment). Finally, when aligning reads from *in vivo* experiments (green alignment), these estimates are used to inject in the alignments the near-exact number of additional reads to compensate for the loss of multimapping reads (purple alignment). (D) Example of correction: FPKM values computed without (light gray) and with the simulation-based correction (dark gray) for 5 paralogous genes of the NOTCH2 family and HES1 as an example of single copy gene.

**Figure S2 (Related to Figure 3).**
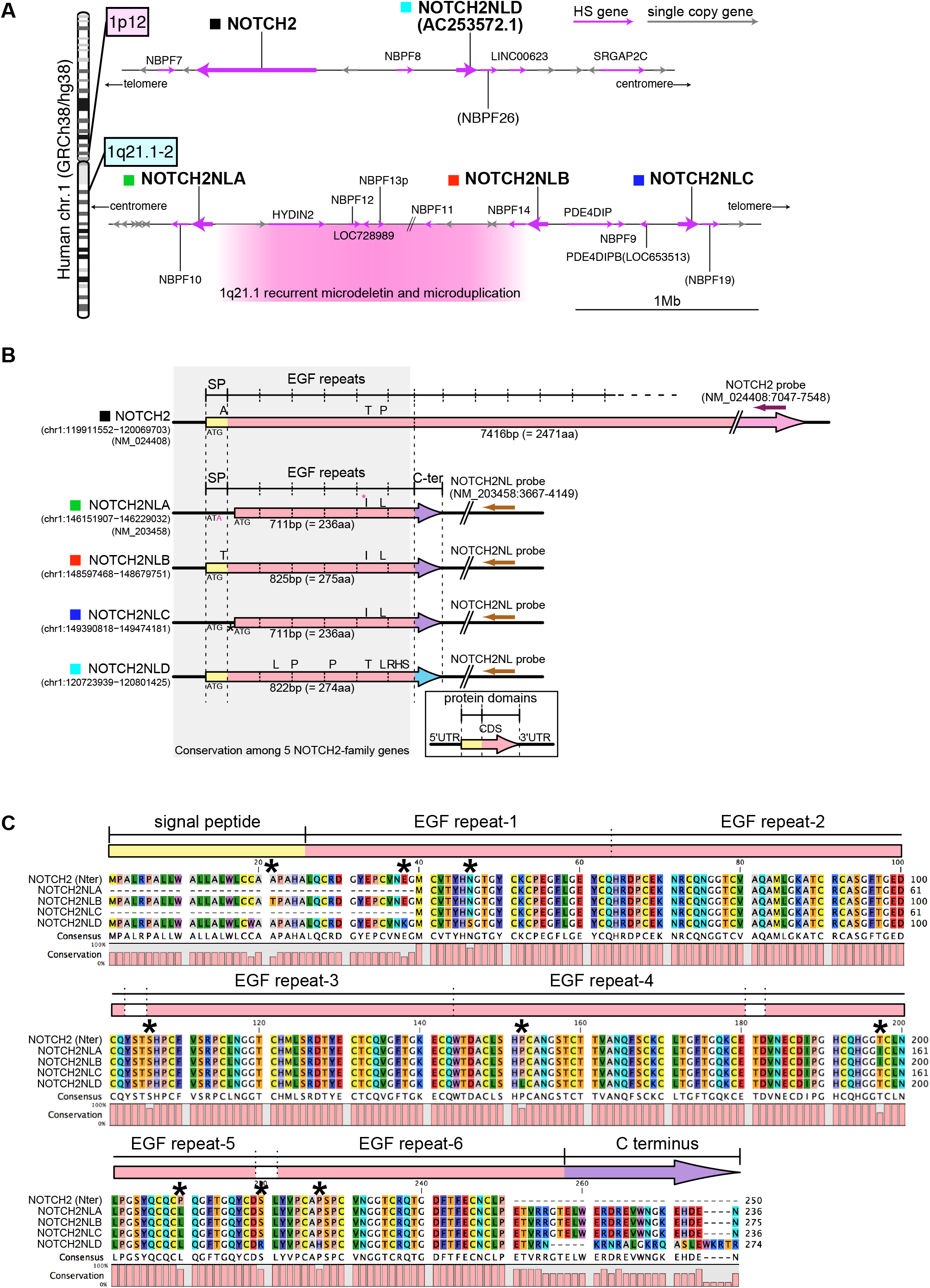
Genomic organization and structure of NOTCH2NL-family gene members. (A) Genomic organization of 1p12 and 1q21.1-2, where NOTCH2-family genes are located. (B) Gene structure of NOTCH2-family members. Protein coding region is indicated by arrows and different colors indicate the protein domains. Amino acid substitutions among the members are indicated above the arrows. (C) Alignment of amino acid sequences of 5 NOTCH2-family gene products. Protein motifs indicated above the alignment is derived from the prediction for NOTCH2 (Uniprot; Q04721). The variable amino acid residues, except for those in the C terminus, are indicated by asterisks.

**Figure S3 (Related to Figure 5).**
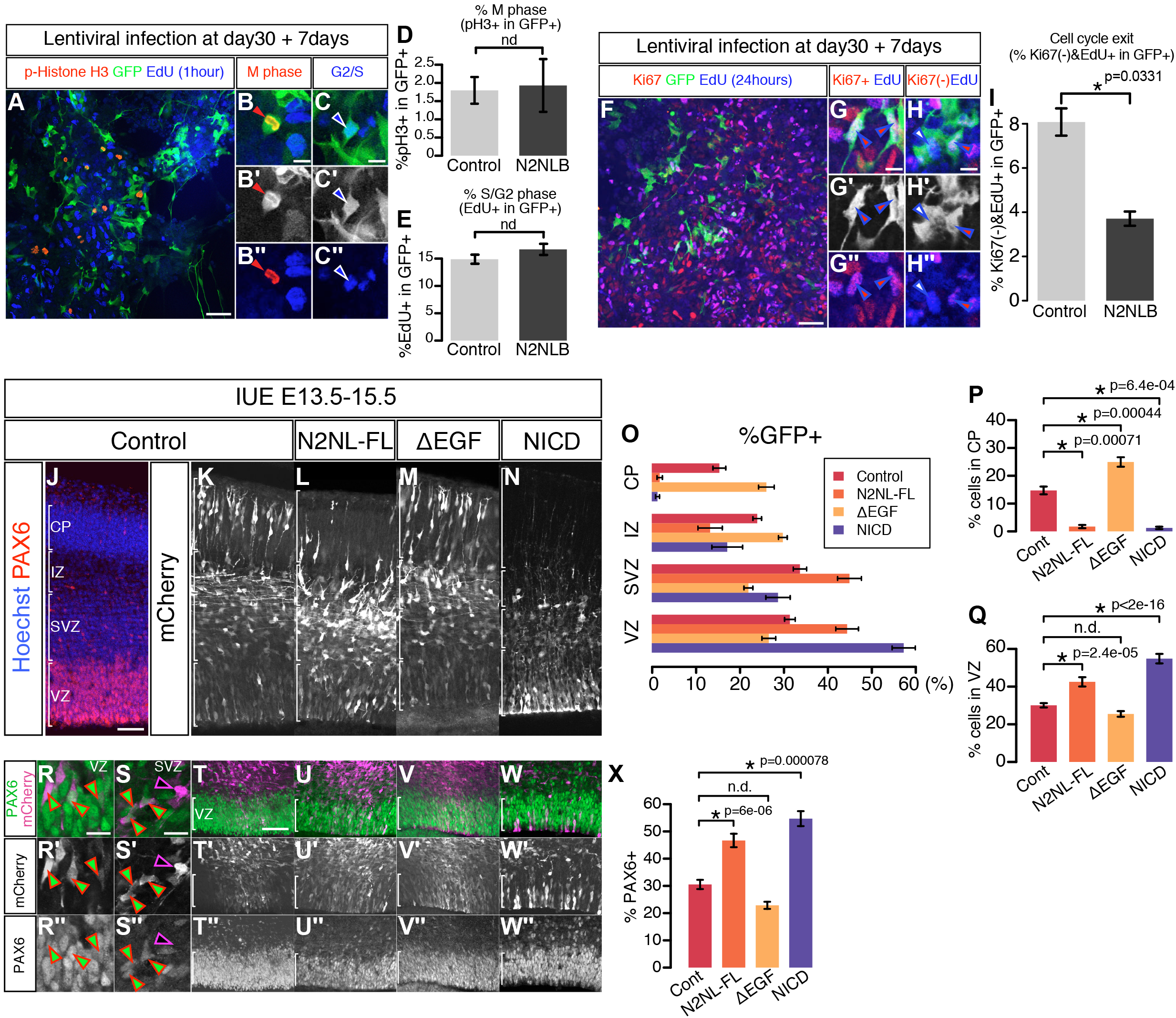
NOTCH2NLB promotes cell cycle reentry and maintenance of cortical progenitors. (A-E) The percentage of mitotic cells and the cells in G2/S phase are quantified using anti-phospho Histone H3 and EdU labelling.(F-I) Cell cycle exit in the human cortical progenitors derived from ESC; NOTCH2NLB introduced by lentiviral infection at day 30 of differentiation, EdU incorporated 24 hours before fixation at day 7 of overexpression. (J,X) *In utero* electroporation of NOTCH2NLB full length (N2NL-FL), NOTCH2NL-EGF repeats deletion (ΔEGF) and mouse NOTCH1 intracellular domain (NICD) in E13.5 mouse cortex, followed by analysis at E15.5. (O-Q) Bin analysis of fractions of mCherry+ electroporated cells in four regions, the CP, IZ, SVZ and VZ., and highlight on VZ and CP (P,Q). Data are represented as mean±sem and p values by Student’s t-test. PAX6 immunoreactivility is examined to quantify the proportion of apical neural progenitors in these four conditions (R-W). (R-X) Proportion of PAX6-positive cells among electroporated cells in the same four conditions. Data are represented as mean±sem and p values by Student’s t-test (D, E and I) one-way ANOVA and bonferroni post hoc test (P, Q. and X). Scale bars; 100μm (A, F, J and T) and 20μm (B, C, G, H, R and S).

**Figure S4 (Related to Figure 6).**
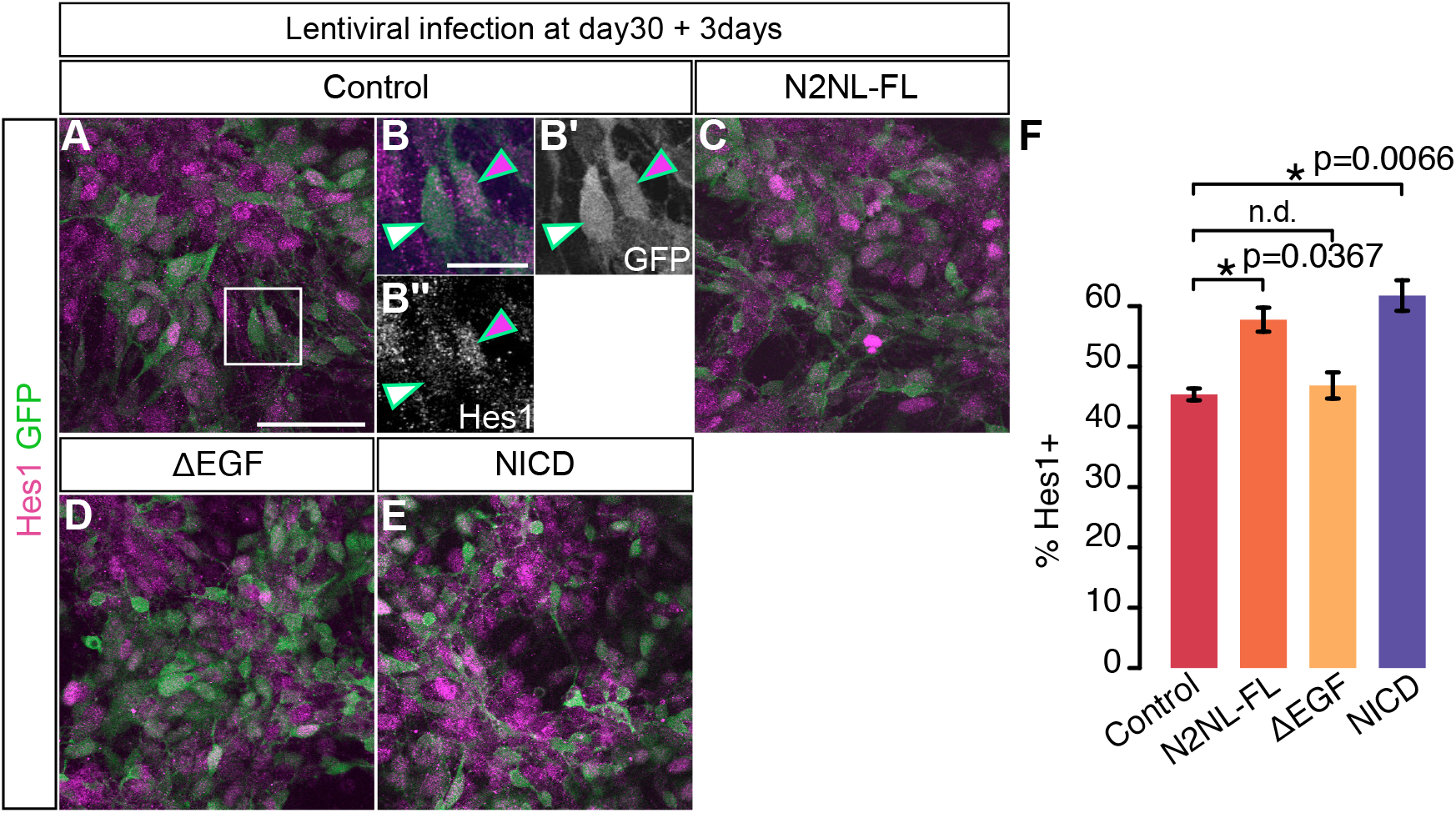
NOTCH2NL activates NOTCH signalling in human cortical progenitors *in vitro*. NOTCH signaling activity was examined using Hes1 as a positive readout of NOTCH signaling in human cortical cells, 3 days following lentiviral infection with control vector (GFP only) or leading to overexpression of NOTCH2NL full length (N2NL-FL), NOTCH2NL-EGF repeats deletion (ΔEGF) or NICD. (F) Quantification of Hes1-expressing cells among GFP labeled cells in each condition. Data are represented as mean±sem and p values by one-way ANOVA and bonferroni post hoc test. Scale bars; 100μm (A) and 20μm (B).

**Figure S5 (Related to Figure 7).**
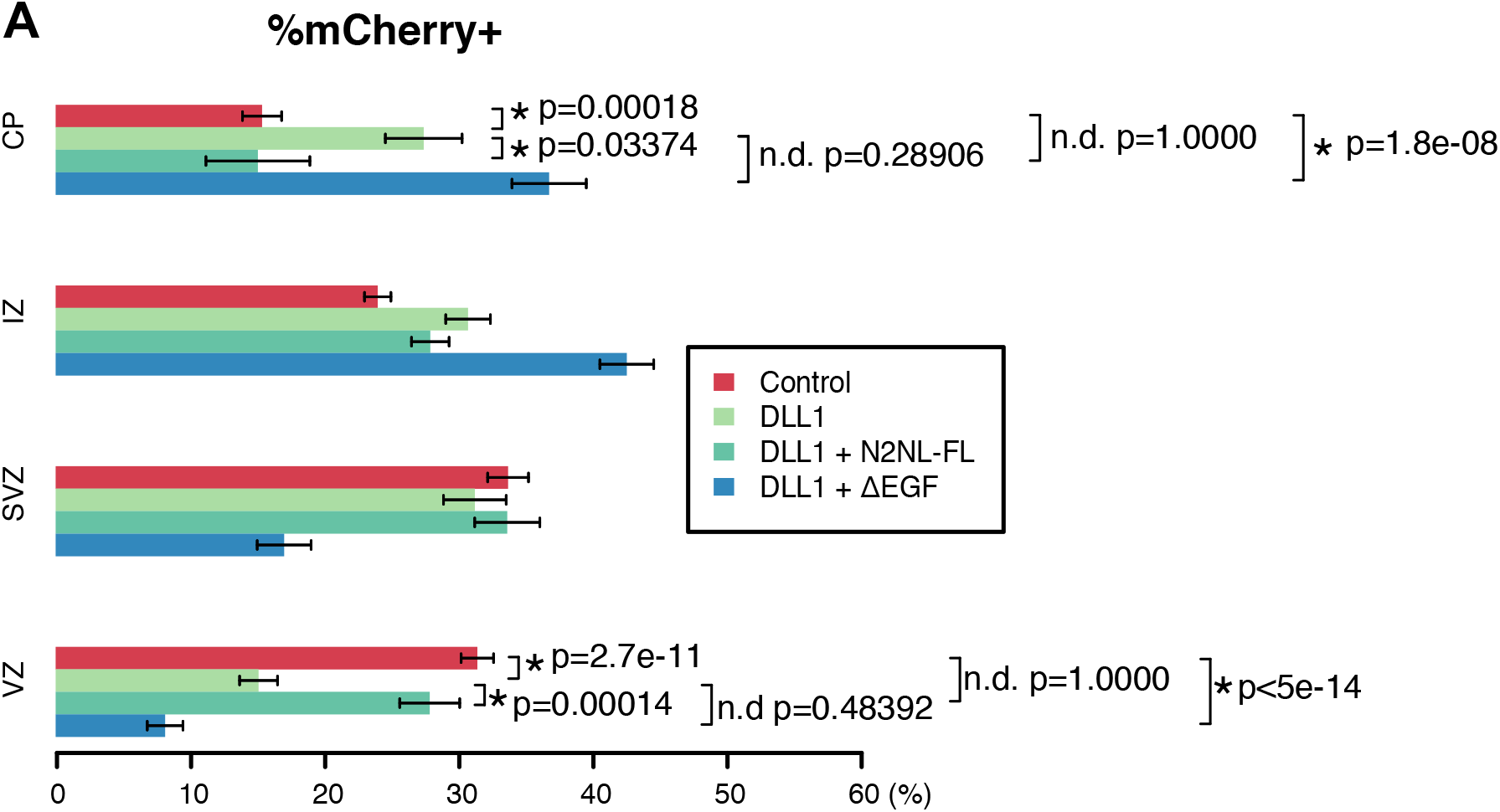
Functional interaction of DLL1 and NOTCH2NL during mouse corticogenesis *in vivo*. Bin analysis of the regional distribution of electroporated cells in the mouse cortex 2 days after *in utero* electroporation at E13.5 (A). Four conditions, Control mCherry alone, DLL1, DLL1 + NOTCH2NLB full length (DLL1+N2NL-FL), DLL1 + NOTCH2NL EGF repeats deletion (DLL1+ΔEGF), were tested. Data are represented as mean±sem and p values by one-way ANOVA and bonferroni post hoc test. the ventricular zone (VZ), the subventricular zone (SVZ), the intermediate zone (IZ), and the cortical plate (CP).

## STAR+METHODS

### CONTACT FOR REAGENT AND RESOURCE SHARING

Further information and requests for resources and reagents should be directed to the Lead Contact Pierre Vanderhaeghen.

## EXPERIMENTAL MODEL AND SUBJECT DETAILS

### Human fetal tissue collection and preparation

The study was approved by three relevant Ethics Committees (Erasme Hospital, Université Libre de Bruxelles, and Belgian National Fund for Scientific Research FRS/FNRS) on research involving human subjects. Written informed consent was given by the parents in each case.

Human fetuses were obtained following medical pregnancy termination. Fetuses aged 7, 9, 12, 15, 17, 19, and 21 GW were used for the RNA sequencing and *in situ* hybridization of cortical tissue. All cases were examined with standard feto-pathological procedures and none displayed clinical or neuropathological evidence of brain malformation. As soon as possible after expulsion (less than 6 hours), the brain was removed using the standard fetal autopsy procedure (Valdés-Dapena and Huff, 1983), frozen in in liquid nitrogen for RNA extraction and embedded as a whole in OCT compound (Tissue-Tek Sakura, VWR Cat# 4583), then snap-frozen in a 2-methylbutane on dry ice bath for histological studies. The non-cortical plate and cortical plate regions of the parietal cortex of GW21 human fetal sample are manually dissected from the 4-5 frozen sections.

### Mice

All mouse experiments were performed with the approval of the Université Libre de Bruxelles Committee for animal welfare. The embryos of a mouse strain ICR (CD1, Charles River Laboratory) were used for *in utero* electroporation. The plug date was defined as embryonic day (E)0.5, and the day of birth was defined as P0.

## METHOD DETAILS

### RNA sequencing

Total RNA was prepared from fetal brain tissues using the total RNA kit I (Omega, Cat# R6834-00). Poly-A tailed RNA was selected and converted to cDNA using TruSeq standard mRNA library prep kit (TruSeq stranded mRNA library prep, illumine, Cat# Cat. No. RS-122-2101). cDNA fragments of 350-700bp size were selected for sequencing (Blue pippin, Sage Science). 150bp of cDNA fragments were sequenced from both ends using HiSeq 2500 with Rapid mode v3 (illumina).

### Transcriptome analysis

#### Defining HS duplicated genes

All genes potentially duplicated in the hominid lineage (Sudmant et al., 2010) were first used as queries in BLAST search for NCBI human reference genome and cDNA databases, and BLAT search on UCSC human reference genome (GRCh38/hg38) to find additional possible duplicated sister genes.

#### Annotation of HS duplicated genes

To obtain a curated annotation file, we first downloaded the RefSeq annotation from the UCSC website. As HS duplicated genes are often poorly annotated, we designed a custom perl script to detect the following conflicting annotations:

- same gene identifier appearing at multiple genomic loci
- multiple gene identifiers for the same genomic loci

Conflicting annotations were then manually corrected by comparing, in the UCSC Genome Browser, UCSC genes, Gencode genes and RefSeq genes tracks. This manual curation led to a total of 342 putative HS gene duplications for which expression was estimated by transcriptome sequencing. The corresponding GTF file was used in all subsequent analysis.

#### Expression bias of HS duplicated genes

Prior to gene expression analysis, we estimated how Cufflinks multi-read correction (-u option) was able to deal with human specific duplicated genes, where the fraction of multi-mapper can be excessive. To this end, we simulated RNA-seq reads with a custom perl script. Given a reference genome and gene annotation in GTF format, this script produces both a set of paired fastq files and the matching ‘reference’ SAM alignment in which all read pairs are perfectly and uniquely mapped. Simulated reads were aligned on the human reference genome hg38 with the STAR program. Gene annotation used to build the reference index was the manually curated Refseq genes previously described.

We then compared Cufflinks expression estimates obtained on the reference alignment and the STAR alignment. As expected, Cufflinks estimates were similar for unique genes without a known paralog, but underestimated for HS genes that were closely related.

To accurately quantify HS gene expression, we performed an alignment correction based on HS gene mappability. This methodology was inspired from the mappability tracks available on the UCSC Genome Browser, but applied directly to transcripts instead on whole genome sequence. For each HS gene, we simulated 1000 read pairs and mapped them back on the reference genome with the STAR program. We then counted the number of read pairs that could be uniquely remapped on each of these genes to estimate their mappability.

Mappability values were further used to correct alignment before estimating expression with Cufflinks. First, we used HTSeq-count to count reads uniquely mapped on every HS gene. Based on their respective mappability value, we estimated the number of reads that were lost due to multi-mapper removal and generated a BAM file with synthetic reads in numbers appropriate to compensate for the loss of multi-mappers. STAR alignment was merged with this compensatory alignment using samtools merge, and this corrected alignment was used as input for Cufflinks. The whole methodology is illustrated in Supplementary figure S1.

#### Sofware and parameters

Reads were aligned with the STAR program (version 2.4.0f1):

STAR –genomeDir STAR_index –readFilesIn R1.fastq.gz R2.fastq.gz – readFilesCommand zcat –runThreadN 5 –outFilterMultimapNmax 30 – outFilterIntronMotifs RemoveNoncanonical –outFilterType BySJout – alignSJoverhangMin 10000 –alignSJDBoverhangMin 1 –outFileNamePrefix output – scoreGenomicLengthLog2scale 0 –scoreStitchSJshift 0 –outFilterMultimapScoreRange 0 –sjdbScore 0 –outFilterMatchNminOverLread 0.3 –outFilterScoreMinOverLread 0.3 alignSJoverhangMin was set to 10000 to prevent STAR from inferring novel splice junctions. We noticed that for the specific case of HS genes located close to each other, STAR produces many false splice junctions linking together paralogs of the same family (e.g. multiple genes of the NBPF family). Also, some default parameters were modified to make sure that spliced reads do not receive any bonus in terms of alignment score. This way, multi-mappers can be better identified based on their sequence identity. Multimappers were further removed and only concordant read pairs were kept for further analysis.

Read counts for HS genes were obtained with HTSeq-count (version 0.6.1p1) : htseq-count -f bam -r pos -s reverse -a 0 alignement.bam genes.gtf > count.out.txt

Cufflinks (version 2.2.1) was used for gene expression and invoked with the following command : cufflinks -o output_dir -p 5 -G genes.gtf -u -b hg38.fa –library-type fr-firststrand aln.corrected.bam

### *In situ* hybridization

*In situ* hybridization using digoxigenin (DIG) -labeled RNA probes was performed as described previously using PCR amplified or plasmid templates (Lambert et al., 2011; Lambot et al., 2005). Alternate sections were processed together in order to allow comparison of the obtained staining. Cryosections of human fetal tissue were fixed in 4% paraformaldehyde (PFA) in PBS for 15min at room temperature and washed with PBST (0.1% Tween20 in PBS) three times. Sections were soaked in 6%H2O2 in PBST for 2 minutes at room temperature and washed three times with PBST. Subsequently sections were incubated in 1μg/ml Proteinase K in PBST for 1 minute and the reaction stopped by incubation in 2mg/ml Glycine in PBST. After washing in PBST, they were fixed again in 4% PFA and 0.2% glutaraldehyde in PBST for 15 minutes at room temperature. For prehybridization, sections are incubated in hybridization solution for one hour at 70C°. The hybridization solution contains 50% formamide, 5xSSC pH4.5, 1% SDS, 50 mg/mL yeast tRNA, and 50 mg/mL heparin. 1.5μg/ml RNA probes in the hybridization buffer were applied to sections and incubated for 16-18 hours at 70C°. The next day, sections were washed in solution ‘1’(50% formamide, 5xSSC pH4.5, 1% SDS) for 15 minutes three times at 70C° and then in solution ‘2’ (50% formamide, 2xSSC pH4.5, 0.11% Tween-20) for 1 minute, three times, at 70C°. After TBST (0.1% Tween20 in TBS) washing, sections were blocked in 5% sheep serum in TBST for a hour at room temperature, followed by overnight incubation in 1/2000 anti-DIG antibody (Merck, Cat#000000011093274910) in TBST at 4C°. On the final day, anti-DIG antibody was washed out by TBST and followed by washing with NTMT (Tris pH 9.5 100 mM, NaCl 100 mM, MgCl2 50 mM, 0.1% Tween-20). Signals were revealed by incubation in NBT/BCIP solution (33μl NBT and 33μl BCIP in 5ml NTMT) at room temperature. Once signal intensity reached to the optimal level, reaction was terminated by postfixation in 4% PFA in PBS for 20 minutes at room temperature. Sections were dehydrated with ethanol series (70%, 90% and 100% for 2 minutes for each) and mounted with the mounting reagent (DPX Mounting Media, Merck Cat#100579). Imaging was performed using a <u>Zeiss Axioplan 2</u> and the intensity and contrast of images were modified using Fiji/ImageJ software where necessary. A sense probe was used as a negative control in each case and revealed no specific staining.

### Digoxigenin (DIG)-labeled riboprobe preparation

Partial cDNA fragments of target genes are amplified by PCR using the primers designed carefully to achieve the specificity to desired target paralogs (Table S4) and subcloned into the cloning vectors (Promega, Cat# A1360).

*In vitro* transcription was performed using linearized plasmids as templates, DIG-labelling mix (Roche, Cat#11585550910) and the T3, T7 or Sp6 RNA polymerases (Roche, Cat#11031163001, Cat#10881767001, and Cat#10810274001). Template DNA was degraded using DNase1 (Roche Cat#10104159001) and RNA was precipitated by ethanol precipitation with LiCl. RNA Probes were analyzed on agarose gel to confirm the purity and the size. All the probes used in this study are summarized in Table S4.

### ESC culture and differentiation

Human ESC H9 (Thomson et al., 1998) cells were propagated using standard procedures on mitotically inactivated mouse embryonic fibroblasts (MEF). Briefly, ESC was cultured in the ES medium, which is Knockout DMEM (Thermo Fisher Scientific, Cat#10829018,) supplemented with 20% Knockout Serum Replacement (Thermo Fisher Scientific, Cat#10828028), 1X Non-essential Amino Acids (Thermo Fisher Scientific, Cat#11140050), 1X Penicillin/Streptomycin (Thermo Fisher Scientific, Cat#15070063), 1X 2-Mercaptoenthanol (Merck, Cat#M6250), 2mM L-glutamine (Thermo Fisher Scientific, Cat#25030081) Medium was changed every day and the cells passaged every 3-4 days using PluriSTEM Dispase-II Solution (Merck, Cat# SCM133). Cortical differentiation from human ESC was performed as described previously (Espuny-camacho et al., 2013). On day 2, cells were dissociated using Stem-Pro Accutase (Thermo Fisher Scientific, Cat#A1110501) and plated on matrigel (hES qualified matrigel BD, Cat#354277) coated dishes at low confluency (5,000–10,000 cells/cm2) in MEF-conditioned hES medium supplemented with 10 mM ROCK inhibitor (Y-27632; Merck, Cat#688000). On day 0 of the differentiation, the medium was changed to DDM (Gaspard et al., 2008), supplemented with B27 devoid of Vitamin A (Thermo Fisher Scientific, Cat#12587010) and 100 ng/ml Noggin (R&D systems, Cat#1967-NG), and the medium was replenished every 2 days. After day 16 of differentiation, the medium was changed to DDM, supplemented with B27 (DDM/B27), and changed every 2 days. At day 24, the progenitors were dissociated using Accutase and cells were resuspended in DDM supplemented with B27 and ROCK inhibitor (10 mM) and plated onto matrigel coated coverslips. Cells are amplified until day 30 in DDM supplemented with B27 for overexpression analysis.

### Lentiviral preparation

HEK293T cells were transfected by packaging plasmids, psPAX2 (Addgene Cat#12260) and pMD2.G (Addgene Cat#12259), and a plasmid of gene of interest in lentiviral backbone (pLenti-CAG-mCherry, pLenti-CAG-NOTCh2NLB-ires-EGFP, pLenti-CAG-NOTCH2NL-ΔEGF-ires-EGFP, pLenti-CAG-NOTCH2NL-ΔC-ires-EGFP and pLenti-CAG-NICD-ires-EGFP). 2 days after transfection, culture medium was collected and viral particles were enriched by filter device (Amicon Ultra-15 Centrifuge Filters, Merck, Cat#<u>UFC910008</u>). Titer check was performed on HEK293T cell culture for every batch of lentiviral preparation.

### Clonal analyses

Clonal analysis was performed as described in the previous study with some modification (Otani et al., 2016). Cortical cells derived from human ESCs were grown on Matrigel-coated coverslips in DDM supplemented with B27 until day 28 in 24 well plate and the coverslips are transferred to 6 well plate (3 coverslips per well). At day 30, the mixture of viruses of Lenti-CAG-mCherry and Lenti-CAG-NOTCH2NLB-ires-EGFP was applied to cells at optimized concentration (which is defined every batch of viral preparation). The medium was changed to fresh DDM supplemented with B27 to wash lentiviruses out one day after intitial infection and the medium subsequently changed every 2-3 days. Each coverslip in a culture well was fixed and immunostained at day 35 (5 days of lentiviral overexpression), day 40 (10 days of lentiviral overexpression) or day 50 (20 days of lentiviral overexpression). The cells with lentiviral overexpression were detected by anti-mCherry and anti-GFP antibodies and the cell types were determined by anti-SOX2 and anti-βIII tubulin antibodies for the cortical progenitors and the differentiated neurons, respectively. The clonal size was defined as a number of cells per cluster, in which cells are located in close proximity each other and are spatially isolated from the neighboring cell clusters in a distance longer than 100μm.

### Acute overexpression in human cortical cells

Lentiviral constructs were used to infect day 30 cortical cells differentiated from human ESC. Cortical cells are prepared as described in the section of clonal analysis. One day after infection, culture medium was changed to wash viruses out. Phenotypes were analyzed 3 and 7 days after infection for Hes1 immunoreactivility and neurogenesis assay by PAX6, SOX2 and βIII tubulin antibodies, respectively.

### Cell cycle labeling assay

A nucleotide analog 5-ethynyl-2’-deoxyuridine (EdU; Merck, Cat#T511285) was incorporated to human cortical cells 24 hours before fixation for cell cycle exit analysis and 1 hour before fixation for G2/S phase labeling. Detection of EdU was performed using Click-iT EdU Alexa Fluor 488/647 Imaging Kit (Thermo Fisher Scientific, Cat# C10337 and Cat#C10340,). For the cell cycle exit analysis, the percentage of cells with EdU-positive and a proliferative cell marker Ki67-negative was examined.

### *In utero* electroporation

*in utero* electroporation was performed as described previously (Dimidschstein et al., 2013; Tiberi et al., 2012a). Briefly, timed-pregnant mice were anesthesized with a mixture of ketamine (Ketalar, 50mg/ml solution injectable, Pfizer) and xylazine (Rompun, 2% solution injectable, Bayer) at E13.5, and each uterus was exposed under sterile conditions. Plasmid solutions containing 1-1.5 mg/ml of DNA were injected into the lateral ventricles of the embryos using a heat-pulled capillary. Electroporation was performed using tweezer electrodes (Nepa Gene, Cat#CUY650P5) connected to a BTX830 electroporator (five pulses of 25 V for 100 ms with an interval of 1s). Embryos were placed back into the abdominal cavity, and mice were sutured and placed on a heating plate until recovery.

### Constructs

Coding sequence of NOTCH2NLB was amplified by PCR from the cDNA library derived from GW9 human fetal cortex using the primers designed on the basis of the sequence of reference genome. The size of PCR fragment was confirmed as expected and PCR fragment was subcloned into the multicloning site before myc tag of a CAG promoter driven expression plasmid (pCAG-IRES-GFP (Dimidschstein et al., 2013; Tiberi et al., 2012a)) by In Fusion cloning (Clontech, Cat#638909). DNA fragment of CAG-NOTCH2NLB-ires-EGFP was transferred to lentiviral plasmid backbone (gift from Cecile Charrier) by restriction digestion and ligation to obtain a lentiviral overexpression construct of NOTCH2NLB (pLenti-CAG-NOTCH2NLB-myc-ires-EGFP). NOTCH2NL deletion constructs (NOTCH2NL-ΔEGF repeats and NOTCH2NL-ΔC terminus) were prepared by PCR amplification using the primers recognizing the desired part of NOTCH2NLB and insertion into the lentiviral vector by In Fusion cloning (pLenti-CAG-NOTCH2NL-ΔEGF-myc-ires-EGFP and pLenti-CAG-NOTCH2NL-ΔC terminus-myc-ires-EGFP). These lentiviral NOTCH2NL plasmids were further modified by replacing EGFP to mCherry for some experiments. Retroviral overexpression construct of rat Delta like-1 (pMX-rDll1-iG) was a gift from Dr. Gotoh (Kawaguchi et al., 2008). The coding sequence of DLL1 C terminally tageed with HA was amplified by PCR to obtain DNA fragment of DLL1-HA and inserted into lentiviral CAG promoter driven overexpression plasmid (pLenti-CAG-DLL1-HA). CBFRE-EGFP was originally developed in a previous study (Mizutani et al., 2007) and obtained from Addgene (#17705).

### Immunofluorescence staining

Mouse embryos were collected 2 days after electroporation and perfused transcardiacally with ice-cold 4% PFA in PBS. Brains were dissected and soaked in 4% PFA solution overnight at 4°C and then sectioned in 100μm thickness using vibrosector (Leica, Cat#VT1000S). Slices were washed with PBST three times and transferred into the blocking solution, which is PBS containing 0.3% Triton X-100 and 3% horse serum, and incubate for 1 hour. Brain slices were incubated overnight at 4°C with the primary antibodies. After three PBST washes, slices were incubated in PBST during 2 hours at room temperature with the secondary antibodies. After washing in PBST, brain sections are amounted on a slide glass with the mounting reagent (DAKO glycerol mounting medium, Cat#C0563). For the human cortical cells, the cells on coverslips were fixed in ice cold 4% PFA in PBS for 15 minutes at room temperature, washed three times with PBST and blocked for 1 hour in the same blocking solution used for mouse brain sections. Then coverslips were incubated in the blocking solution containing primary antibodies overnight at 4°C. Coverslips were washed three times in PBST and incubated in PBST containing the secondary antibodies. Coverslips were mounted on a slide glass with the mounting reagent. Imaging was performed using a confocal microscope Zeiss LSM780 and Images are processed by Fiji/ImageJ software.

### Detection of DLL1 protein on the plasma membrane

Engineered CHO cell lines, which expresses Delta like-1 C-terminally fused with mCherry under the tetracyclin-dependent promoter (Sprinzak et al., 2010), were used for this analysis. Experimental scheme was modified from the preceding studies (LeBon et al., 2014; Sprinzak et al., 2010). Cells were transfected with the overexpression plasmid of NOTCH2NLB or NOTCH2NL deletion mutant of EGF repeats, both of which are C-terminally fused with myc epitope tag, using the X-tremeGENE HP DNA Transfection Reagent (Merck, Cat#6366244001). Then cells were passaged and grown in low confluency (20,000 cells/cm2) in CHO cell medium (alpha MEM supplemented with 10% FBS, 1X Penicillin/Streptomycin, and L-Glutamine) containing 100ng/ml Doxycycline hydrochloride (Merck, D3447). Recombinant chimeric protein of Notch1 extracellular domain fused with mouse Fc fragment of IgG (Notch1-Fc, R&D systems, Cat#5267-TK-050) was applied to the culture medium and incubated on ice for 1 hour. Cells were acutely fixed in 4% PFA in PBS for 15 minutes on ice and washed with PBST three times. Then cells were permeabilized and blocked in the blocking solution containing 0.3% Triton X-100, and followed by the incubation in the blocking solution containing primary antibodies overnight; anti-DLL1 (rabbit, Santa Cruz, Cat#sc9102) and anti-Myc (goat, abcam, Cat#ab9132) antibodies. On the next day, cells were washed and labeled with the secondary antibodies; anti-Mouse IgG antibody conjugated with Alexa488 for the detection of Notch1-Fc, anti-Rabbit IgG antibody conjugated with Alexa405 for the detection of DLL1 and anti-Goat IgG antibody conjugated with Alexa647 for the detection of Myc. Images containing three channels were obtained using a confocal microscopy Zeiss LSM780. Myc-positive NOTCH2NL-expressing cells and Myc-negative control cells were imaged in whole Z axis with the interval of 0.6μm.

### Confocal microscopy

Confocal images were obtained with Zeiss LSM 780 driven by ZEN 2012 software and equipped with 10x 0.30, 20x 0.8 and 40x 1.1W objectives. Microscope disposed of argon, helium-neon and 405 nm diode lasers.

## QUANTIFICATION AND STATISTICAL ANALYSIS

### Statistical analysis

Data in figure panels reflect 3 or more independent experiments performed on different days. An estimate of variation within each group of data is indicated using standard error of the mean (SEM).

We performed unpaired Student’s t-test for assessing the significance of differences in the analyses containing two conditions and one-way ANOVA and bonferroni post hoc test in the analyses containing more than three conditions using R software (See each figure for details).

### *In utero* electroporation in mouse

For quantification of the results of *in utero* electroporation experiments, we obtained the image of whole cortical thickeness in the columnar imaging window with a 300μm width at the center of electroporation. In every experiment, we obtained multichannel images of fluorescent marker EGFP or mCherry to identify electroporated cells and immunostaining of one or two antibodies in addition to Hoechst33342 counter staining. We counted the numbers of electroporated cells by fluorescent markers EGFP or mCherry in the cortical regions from the most apical ventricular zone to the most basal cortical plate (CP). For E15.5 and E16.5 samples, four regions (VZ, SVZ, IZ and CP) were determined by the cellular density revealed by Hoechst 33342 and by the VZ marker PAX6 (VZ; dense/PAX6-positive, SVZ; dense/PAX6-negative, IZ; sparse, and CP; dense). For E18.5 samples, six regions (VZ, SVZ, IZ, SP, lower and upper CP) were determined by cellular density and the PAX6 and MAP2 immunostainings (VZ: dense/PAX6-positive, SVZ; dense/PAX6-negative, IZ; sparse, SP; sparse/MAP2-strongly positive, CP; dense/MAP2-weakly positive). The CP was further subdivided equally into the lower and upper parts. The percentage of electroporated cells in each region is determined as the number of electroporated cells in a given region divided by the total number of electroporated cells in whole cortical wall. The percentage of PAX6-positive cells are defined as the number of PAX6 immunopositive electroporated cells over total number of electroporated cells in whole cortical wall. Similarly, the percentage of NOTCH reporter-positive cells are defined as the number of NOTCH reporter-positive electroporated cells over the total number of electroporated cells in whole cortical wall.

### Clonal assay in human cortical cells

The number of cells with lentiviral overexpression per clonal cluster was counted for 25-30 clusters of each condition (control-mCherry and NOTCH2NLB-ires-EGFP) in an experiment. By repeating and pooling the data of three independent experiments, we obtained the clonal size for 80-100 clones for each tested condition.

### Detection of DLL1 protein on the plasma membrane

Under the culture condition that 2-10 cells are making an isolated cluster with a significant distance from neighboring one, we selected clusters of solely Myc-positive (NOTCH2NLB-overexpression) or Myc-negative cells (non overexpression as control) to avoid the mixed cellular cluster. For each cluster of cells, a series of confocal images were obtained along the Z axis with a 0.6μm interval, in which each image contains three channels of NOTCH1-Fc (Alexa488), DLL1 (Alexa405) and NOTCH2NLB-overexpression by Myc immunofluorescence (Alexa647). The signal intensities of NOTCH1-Fc and DLL1 were measured in every optical section for a cell cluster and pooled to quantify total NOTCH1-Fc signal and total DLL1 signal for a given cluster of cells. These values from 20-30 clusters per experiment were plotted in two-dimensional scatter plot we then drew regression lines for each signals using R. Then the ratio of NOTCH1-Fc signal over DLL1 signal was calculated for all clusters analyzed. Data are represented as mean±sem and Student’s t-test was performed for each of three independent experiments.

## DATA AND SOFTWARE AVAILABILITY

The access number for the data for RNAseq reported in this study is NCBI GEO: xxxxxxxx.

### Supplementary tables

Table S1 Transcriptome of HS genes during human corticogenesis (Related to Figure 1)

Table S2 HS genes identified in the steps of transcriptome screening (Related to Figure 1)

Table S3 Detailed information of probes used for *in situ* hybridization (Related to Figure 2)

